# Immune and malignant cell phenotypes of ovarian cancer are determined by distinct mutational processes

**DOI:** 10.1101/2021.08.24.454519

**Authors:** Ignacio Vázquez-García, Florian Uhlitz, Nicholas Ceglia, Jamie L.P. Lim, Michelle Wu, Neeman Mohibullah, Arvin Eric B. Ruiz, Kevin M. Boehm, Viktoria Bojilova, Christopher J. Fong, Tyler Funnell, Diljot Grewal, Eliyahu Havasov, Samantha Leung, Arfath Pasha, Druv M. Patel, Maryam Pourmaleki, Nicole Rusk, Hongyu Shi, Rami Vanguri, Marc J. Williams, Allen W. Zhang, Vance Broach, Dennis Chi, Arnaud Da Cruz Paula, Ginger J. Gardner, Sarah H. Kim, Matthew Lennon, Kara Long Roche, Yukio Sonoda, Oliver Zivanovic, Ritika Kundra, Agnes Viale, Fatemeh N. Derakhshan, Luke Geneslaw, Ana Maroldi, Rahelly Nunez, Fresia Pareja, Anthe Stylianou, Mahsa Vahdatinia, Yonina Bykov, Rachel N. Grisham, Ying L. Liu, Yulia Lakhman, Ines Nikolovski, Daniel Kelly, Jianjiong Gao, Andrea Schietinger, Travis J. Hollmann, Samuel F. Bakhoum, Robert A. Soslow, Lora H. Ellenson, Nadeem R. Abu-Rustum, Carol Aghajanian, Claire F. Friedman, Andrew McPherson, Britta Weigelt, Dmitriy Zamarin, Sohrab P. Shah

## Abstract

High-grade serous ovarian cancer (HGSOC) is an archetypal cancer of genomic instability patterned by distinct mutational processes, intratumoral heterogeneity and intraperitoneal spread. We investigated determinants of immune recognition and evasion in HGSOC to elucidate co- evolutionary processes underlying malignant progression and tumor immunity. Mutational processes and anatomic sites of tumor foci were key determinants of tumor microenvironment cellular phenotypes, inferred from whole genome sequencing, single-cell RNA sequencing, digital histopathology and multiplexed immunofluorescence of 160 tumor sites from 42 treatment-naive HGSOC patients. Homologous recombination-deficient (HRD)-Dup (*BRCA1* mutant-like) and HRD-Del (*BRCA2* mutant-like) tumors harbored increased neoantigen burden, inflammatory signaling and ongoing immunoediting, reflected in loss of HLA diversity and tumor infiltration with highly-differentiated dysfunctional CD8^+^ T cells. Foldback inversion (FBI, non-HRD) tumors exhibited elevated TGFβ signaling and immune exclusion, with predominantly naive/stem-like and memory T cells. Our findings implicate distinct immune resistance mechanisms across HGSOC subtypes which can inform future immunotherapeutic strategies.

**HIGHLIGHTS:**

- Multi-region, multi-modal profiling of malignant and immune cell phenotypes in ovarian cancer
- Anatomic site specificity is a determinant of cancer cell and intratumoral immune phenotypes
- Tumor mutational processes impact mechanisms of immune control and immune evasion
- Spatial topology of HR-deficient tumors is defined by immune interactions absent from immune inert HR-proficient subtypes

## INTRODUCTION

Genomic instability is a hallmark of human cancer, which often occurs due to impaired DNA repair mechanisms such as homologous recombination (HR), leading to chromosomal copy number alterations and structural genomic rearrangements. The nature of genomic instability has fundamental relevance to cancer etiology and evolution, and anti-tumor immune responses. Increasingly, the role of structural alterations in eliciting immune response and escape has been highlighted ^1,2^. These and other advances have prompted open questions about how anti-tumor adaptive immunity is impacted by specific genomic instability mutational processes, defined by acquired structural variation patterns in cancer genomes ^3–7^.

High-grade serous ovarian cancer (HGSOC) is an archetypal tumor of genomic instability. The principal defining features of HGSOCs are profound structural variations in the form of copy number alterations and genomic rearrangements on a genetic background of near ubiquitous mutations in *TP53,* rendered bi-allelic through loss of heterozygosity of chromosome arm 17p ^8,9^. Somatic and germline alterations in the HR repair pathway genes such as *BRCA1* and *BRCA2* mutations, lead to HR deficiency (HRD) in approximately half of HGSOCs ^10^. Beyond gene alterations, recent work by our group and others has identified distinct patient strata associated with endogenous mutational processes inferred from structural variation patterns in whole genome sequencing. These include HRD subtypes (*BRCA1*-associated tandem duplications: HRD-Dup; *BRCA2*-associated interstitial deletions: HRD-Del), *CCNE1*-amplified associated foldback-inversion (FBI) bearing tumors and *CDK12*-associated tandem duplicators (TD) ^11,12^, amongst related mutational processes defined by copy number alteration ^7^. Notably, the mutational processes are associated with different outcomes, with FBI and TD tumors exhibiting the worst prognosis ^5,11,12^.

Another distinctive property of HGSOC is that patients often present with widespread disease at diagnosis. HGSOC is thought to originate in the fimbriated end of fallopian tube epithelium ^13,14^ and a long latency allows for broad periods of genomic instability, clonal diversification and tumor-immune interactions to unfold in the heterogeneous microenvironments of the peritoneal cavity ^15–17^. Little is known about how the local tissue microenvironment determines anti-tumor immunity. Furthermore, underlying relationships between mutational processes, tumor signaling, microenvironment composition, and immune cell phenotypes remain poorly understood. Accordingly, improved definition and enumeration of the constituent cell types, their derivative cellular phenotypes and their localization both within and between patients are key steps to understanding malignant cell and tumor microenvironment interactions, and ultimately responses to immunotherapeutic interventions. We hypothesized that cellular composition, topology and phenotypic states comprising TMEs differ according to their underlying mutational processes and spatial context. To address this, we designed a prospective multi-modal, multi-site study, capturing mutational processes from whole genome sequencing, disaggregated single cell transcriptome and *in situ* cellular imaging of protein measurements in HGSOC at scale. Our findings reveal that distinct immunostimulatory and immunosuppressive mechanisms co-segregate with both site of disease and underlying mutational process, yielding new insights into malignant-immune microenvironment interactions for cancers of genomic instability with implications for personalized therapeutic strategies.

## RESULTS

### Patient cohort and multi-region, multi-modal profiling

We studied tumors from treatment-naive, newly diagnosed HGSOC patients (**Tab. S1**) consented to a biospecimen banking protocol approved by the institutional review board. We collected multi-site tissue biopsies (n=160) from pre-treatment patients (n=42) undergoing laparoscopy or primary debulking surgeries over a 24-month period. Anatomic site collections included adnexa (ovary and fallopian tube), omentum, peritoneum, bowel, ascites and other intraperitoneal sites (**Fig. 1A**). Clinical characteristics of all patients are summarized in **Fig. 1B** and **Tab. S1**.

**Figure 1.**
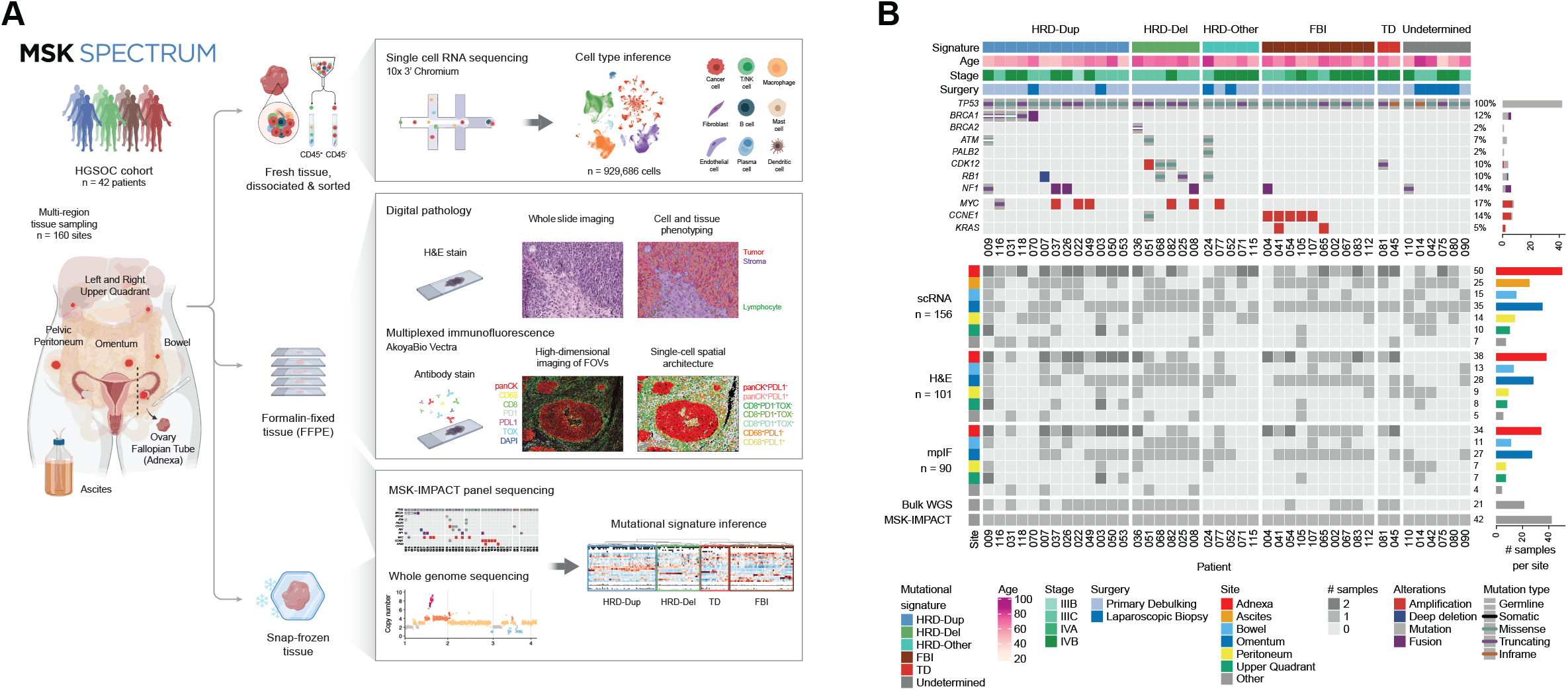
Multi-region, multi-modal profiling of malignant cells and the TME. **A)** Schematic of the MSK SPECTRUM specimen collection workflow including surgery, single-cell suspensions for scRNA-seq and biobanking of snap-frozen and FFPE samples. **B)** Cohort overview. Top panel: Oncoprint of selected somatic and germline mutations per patient and cohort-wide prevalence by MSK-IMPACT. Patient data include mutational signature subtype, patient age, staging following FIGO Ovarian Cancer Staging guidelines, and type of surgical procedure. Bottom panel: Sample and data inventory indicating number of co-registered multi-site datasets: single-cell RNA sequencing, H&E whole-slide images, multiplexed immunofluorescence, whole genome sequencing and targeted panel sequencing (MSK-IMPACT).

We profiled patient samples with five different data modalities (**Fig. 1A**). (1) Fresh tissue samples were collected, dissociated, flow-sorted for live CD45^+^ and CD45^-^ cells to enrich immune cell populations, and processed for transcriptomic profiling using 10x 3’ single-cell RNA sequencing (scRNA-seq) from multiple sites (n=156) of 41 patients, within a one-day workflow from surgery to sequencing library preparation (**Methods**). This yielded a total of 929,686 quality-filtered single cell transcriptomes with an average of ∼23k per patient (∼6k per site), including data from CD45^+^ and CD45^-^ populations (**Tab. S4**). (2) Formalin fixed paraffin embedded tissues (FFPE) were employed for whole-slide hematoxylin and eosin (H&E) staining. For each specimen with scRNA-seq, 101 site-matched H&E sections from 35 patients were digitally scanned and annotated for computational analysis of lymphocytic infiltration. (3) For each specimen with scRNA-seq, site-matched FFPE tissue sections adjacent to the H&E section were stained and imaged by multiplexed immunofluorescence (mpIF) for major cell types and immunoregulatory markers (DAPI, panCK + CK8/18, CD8, CD68, TOX, PD1, PDL1) on the AkoyaBio Vectra platform (**Methods**). A total of 10,663,919 cells from 1,194 quality-filtered fields of view across 90 tissue samples from 35 patients were identified for downstream spatial topology analysis. (4) FDA-approved clinical sequencing of 468 cancer genes (MSK-IMPACT) was obtained on DNA extracted from FFPE tumor and matched normal blood specimens to establish mutational status of known high prevalence alterations in HGSOC: *TP53* (100%)*, BRCA1* (12%)*, BRCA2* (2%)*, CCNE1* high level amplification (12%)*, MYC* high level amplification (17%)*, CDK12* (10%), and *RB1* (10%) (**Fig. 1B**), comprising a representative set of the cohort profiled by MSK-IMPACT (**Fig. S1**). (5) Lastly, where available, snap-frozen tissues were processed to obtain matched tumor-normal whole genome sequencing (WGS) on a single representative site of the subset of 41 patients with scRNA-seq to derive mutational processes from single nucleotide and structural variant mutational signatures. We assigned established mutational signatures by WGS, yielding 13 HRD-Dup, 6 HRD-Del, 10 FBI and 2 TD patients, as well as 5 additional HRD cases labelled HRD-Other, which we identified based on Myriad HRD testing or the presence of inactivating mutations in HR genes detected by MSK-IMPACT (**Fig. 1B**, **Tab. S1**, **Tab. S2**, **Methods**).

### Cellular constituents of the HGSOC tumor microenvironment vary by patient and site

We first constructed a cell map from the scRNA-seq data, organized into nine broad cellular lineages: 251,837 epithelial cells, 289,952 lymphoid cells (T cells, B cells, plasma cells, NK cells), 207,288 myeloid cells (monocytes/macrophages, dendritic cells, mast cells) and 180,609 stromal cells (fibroblasts, endothelial cells) (**Fig. 2A,B**). Non-malignant cells were well separated by cell type in UMAP space with cells from different patients intermixed. In contrast, ovarian cancer cells were mainly separated by patient, quantified through patient specificity scores with a shared nearest neighbor graph (SNN) (**Methods**). We attributed the high degree of patient specificity of ovarian cancer cells (**Fig. 2A,C**) to tumor cell-specific somatic copy number alterations driving concomitant gene dosage effects. Fibroblasts and myeloid cells also exhibited patient specificity, which we interpreted as unique tumor-associated responses (**Fig. 2C**). Notably, patient variation in malignant cells was not attributed to previously reported bulk expression signatures ^18,19^ (**Fig. 2D**). Cancer cells were primarily classified into the proliferative (19.5%) and differentiated (78.8%) subtypes (**Fig. 2D**), depending on whether they were cycling or not. Myeloid cells were most commonly classified into the immunoreactive subtype (76.6%). None of the other immune cell types contributed to the immunoreactive subtype and were mainly assigned to the proliferative (4%-23.2%) or differentiated (62.9%-87.1%) subtypes (**Fig. 2D**). Endothelial cells were likewise comprised of cells classified as proliferative (23.7%) or differentiated (69.8%). However, fibroblasts were the only cell type to show enrichment for mesenchymal classification (62.6%). We thus concluded that gene expression signatures defining previously reported transcriptional subtypes of HGSOC predominantly reflect cell type composition, rather than intrinsic variation in malignant cell phenotypes.

**Figure 2.**
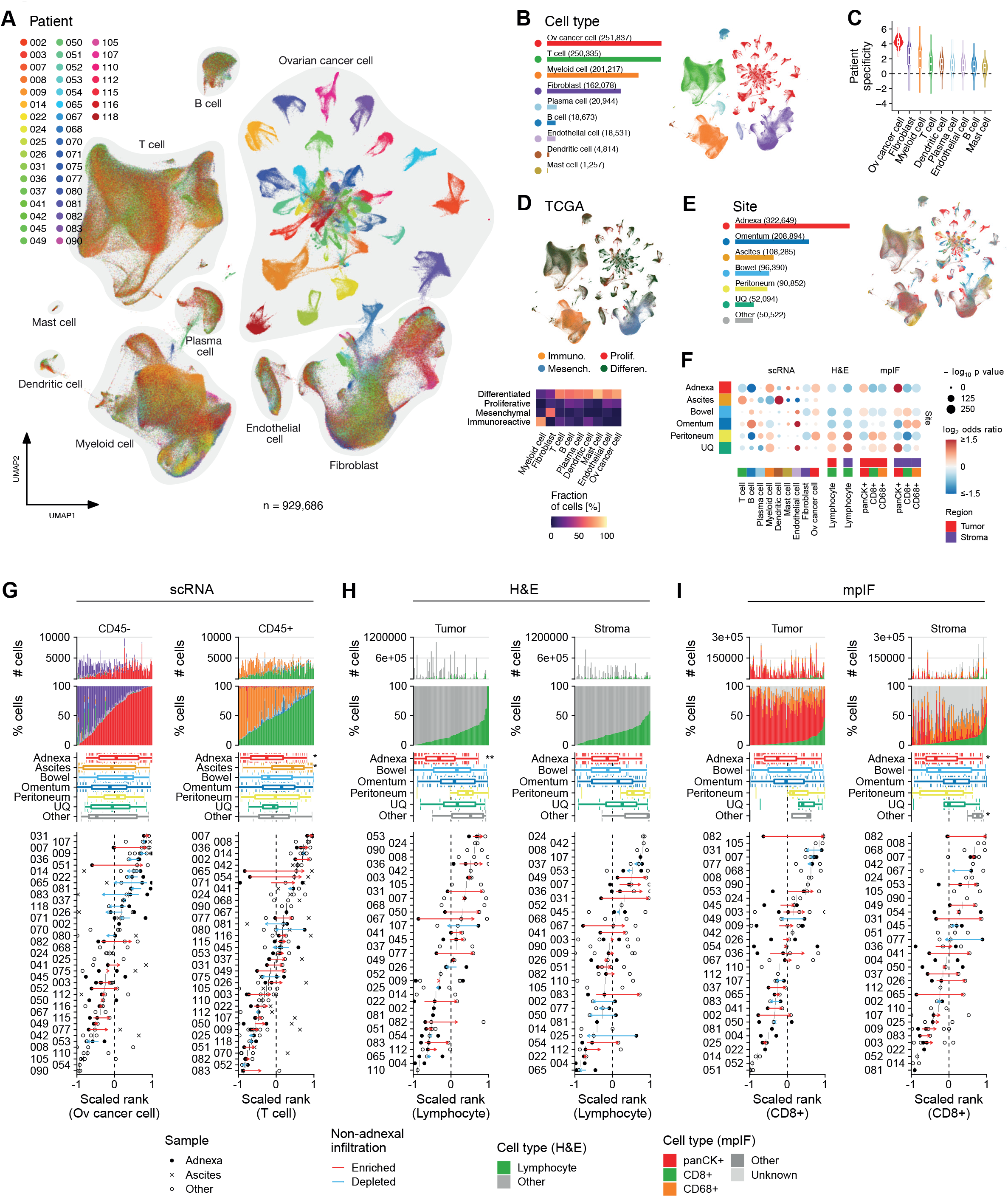
The site-specific tumor microenvironment of HGSOC at single-cell resolution. **A)** UMAP of cells profiled by scRNA-seq colored by patient. Cell types as defined by CellAssign ^67^ are highlighted with grey outlines. **B)** Number of cells identified per cell type next to UMAP colored by cell type. **C)** Patient specificity per cell type computed as the number of observed neighboring cells coming from the same patient over the number of expected neighboring cells coming from the same patient with zero indicating a uniform distribution. **D)** Upper: UMAP colored by TCGA transcriptional subtype. Lower: Fraction of cells assigned to a given TCGA transcriptional subtype per cell type. Columns in the heatmap add up to 100%. **E)** Number of cells profiled per tumor site next to UMAP colored by tumor site. **F)** Site-specific enrichment of cell type composition in scRNA, H&E and mpIF data fitted using a generalized linear model (GLM). GLMs for H&E and mpIF data are separated by tumor and stroma regions. Color gradient indicates log_2_ odds ratios (enrichment: red, depletion: blue) and sizes indicate the BH-corrected -log_10_(p value). **G)** Cell type composition based on scRNA for CD45^-^ samples (left) and CD45^+^ samples (right). Upper panels: Absolute and relative cell type numbers. Middle panels: Box plot distributions of sample ranks with respect to tumor site. Lower panels: Dot plot of sample ranks grouped by patient. Colored arrows indicate enrichment (red) or depletion (blue) of ovarian cancer cells (left) and T cells (right) in non-adnexal over adnexal samples. **H)** Cell type composition based on H&E with lymphocyte ranks in tumor-rich (left) and stroma-rich (right) compartments. Panels analogous to Fig. 2G. **I)** Cell type composition based on mpIF with CD8^+^ T cell ranks in tumor-rich (left) and stroma-rich (right) compartments. Panels analogous to Fig. 2G. \**P*<0.05, \*\**P*<0.01.

We next analyzed cell type composition variation with respect to anatomic sites within patients, ranging between 322,649 cells derived from adnexal samples to 52,094 from the upper quadrants (**Fig. 2E**). We compared the relative proportions of cell types in the adnexa (i.e. potential primary lesions of the fallopian tube and ovary), ascites, and distal sites throughout the peritoneal cavity. CD45^+^ samples ranged from myeloid-rich to lymphoid-rich within and between patients, with adnexal samples significantly depleted for T cells (Mann-Whitney U test, BH-corrected, q-value = 0.0195), B cells (q-value = 0.0041) and dendritic cells (q-value = 0.0060), while in contrast, ascites samples were enriched for T cells (q-value = 0.0195) and dendritic cells (q-value < 0.0001). Notably, T cell fractions in distant intraperitoneal sites were consistently higher than in adnexal samples (22 of 32 patients) (**Fig. 2F,G**). This was corroborated with site-matched whole-slide H&E analyses, where intratumoral lymphocyte proportions were increased in distant non-adnexal sites compared with paired adnexal sites in 17 out of 22 patients (**Fig. 2F,H**), and enriched for intratumoral CD8^+^ T cells in paired non-adnexal over adnexal samples in 14 out of 21 patients (**Fig. 2F,I**) from site-matched mpIF.

### Phenotypic differentiation of T cells exhibits inter-site heterogeneity

We next assessed variation of constituent T cell subtypes and functional states at distinct tumor sites. We identified 41 distinct T and NK cell clusters, broadly defining CD4^+^ T cells (clusters 1-14), CD8^+^ T cells (clusters 15-22), γδ T cells (cluster 23), NK cells (clusters 24-33) and cycling cells (clusters 34-41) (**Fig. 3A-B**; **Fig. S3A,C**). We found a graded enrichment and depletion of specific T and NK cell clusters across UMAP space (**Fig. 3A**, bottom) in adnexal and non-adnexal sites. Generalized linear modeling (GLM) revealed pronounced site-specific differences in cluster composition, with the biggest differences between adnexal and ascites samples (**Fig. 3B**, **Methods**). In particular, CD4^+^ naive/stem-like and central memory T cells (clusters 1-2) were depleted in the adnexal samples but were enriched in ascites samples (**Fig. S3B**, log_2_ odds_CD4.T.naive.mem|adnexa_ = -0.49, log_2_ odds_CD4.T.naive.mem|ascites_ = +0.91). Conversely, dysfunctional CD4^+^ and CD8^+^ T cells (clusters 4-6 and 18-20) were depleted in ascites samples (log_2_ odds_CD4.T.dys|ascites_ = -0.62, log_2_ odds_CD8.T.dys|ascites_ = -0.85) but enriched in adnexal samples. This is consistent with chronic antigen exposure leading to dysfunction at higher rates in the adnexa (log_2_ odds_CD4.T.dys|adnexa_ = +0.12, log_2_ odds_CD8.T.dys|adnexa_ = +0.37). Regulatory T cell (8-10) and regulatory NK cell clusters (27-33) were likewise enriched in the adnexal samples, suggesting that they may carry out immunomodulatory feedback at these sites (**Fig. 3B**). Notably, when ranked by relative fraction of naive/stem-like and central memory T cells or dysfunctional T cells, we observed a high inter- and intra-patient variability in the relative composition of the T cell clusters (**Fig. 3C**, top). Distributions of scaled ranks reflected a marked inter-site variation in tumor infiltration (**Fig. 3C**, box plot panel). Within patients, adnexal samples were depleted for naive/stem-like and central memory T cells in 26 out of 32 patients and enriched for dysfunctional T cells in 23 out of 32 patients.

**Figure 3.**
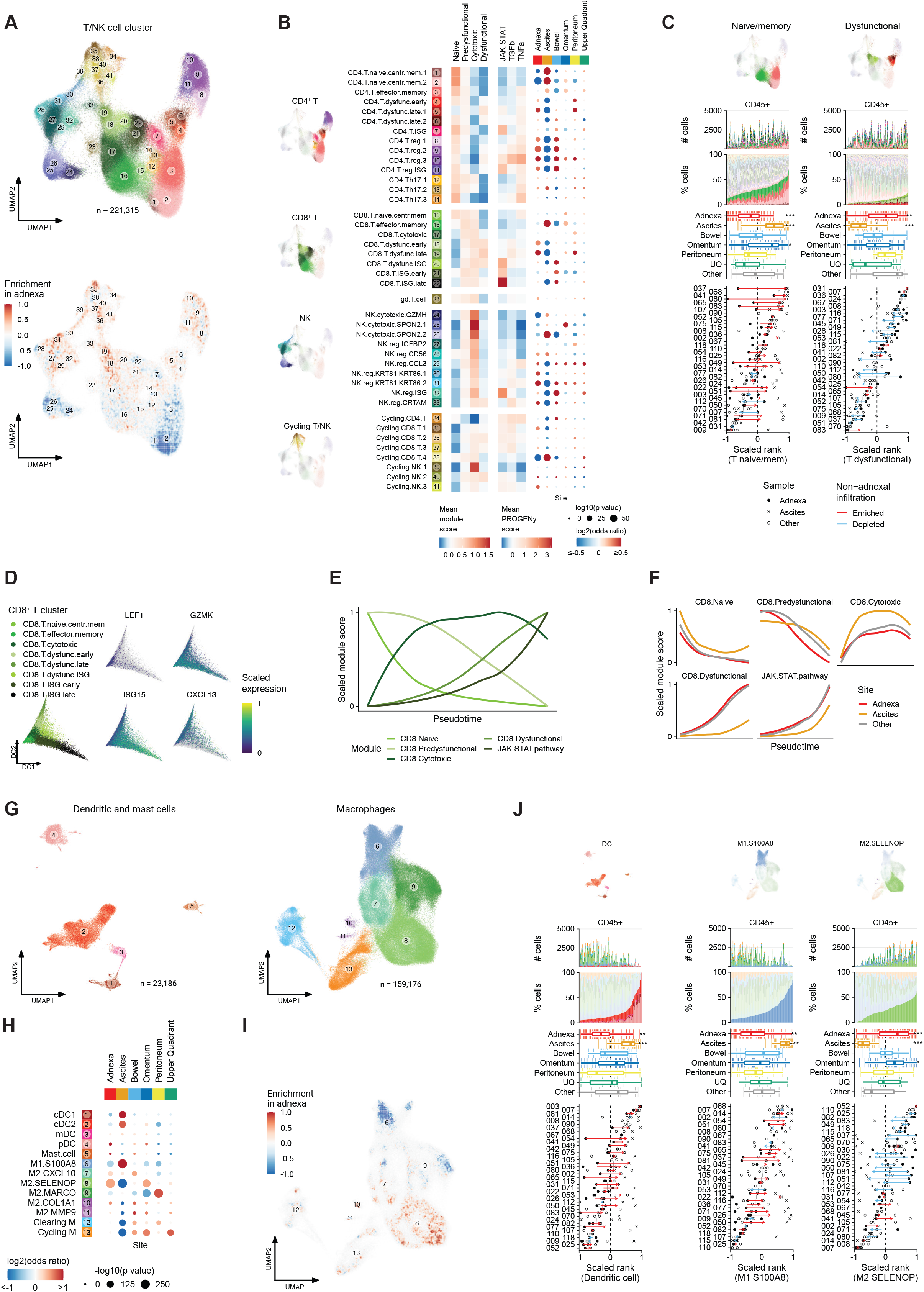
Adnexal samples exhibit increased T cell dysfunction and are enriched for immunosuppressive macrophages. **A)** Upper: UMAP of T and NK cell clusters profiled by scRNA-seq. Clusters are colored and numbered to reference cluster labels in B). Lower: Kernel density estimates in UMAP space for adnexal-enriched (red) over non-adnexal enriched (blue) clusters. **B)** Heatmap of PROGENy signaling pathway activity scores (left) and T cell state module scores (middle) across CD4^+^ T, CD8^+^ T, γδT, NK and Cycling cell clusters. Dot plot panel (right) shows site-specific enrichment of T/NK cell clusters using GLM. Color gradient indicates log_2_ odds ratios (enrichment: red, depletion: blue) and sizes indicate the BH-corrected -log_10_(p value). **C)** T/NK cell cluster composition based on scRNA ranked by fraction of T naive/memory clusters (left) or fraction of T dysfunctional clusters (right). Panels analogous to Fig. 2G-I. **D)** Left: Diffusion maps of the subset of CD8^+^ T cells profiled by scRNA-seq, colored by CD8^+^ T cell cluster. Right: Relative expression of genes marking CD8^+^ T cell clusters in diffusion space. **E)** Scaled module scores with respect to pseudotime inferred from diffusion components. **F)** Analogous to E), separated out by tumor site. **G)** UMAPs of dendritic cells and mast cells (left) and macrophages (right) profiled by scRNA-seq, colored and numbered by cluster as in H). **H)** Site-specific enrichment of myeloid cell clusters using GLM, analogous to B). **I)** Differences in kernel density estimates in UMAP space for macrophages in adnexal enriched (red) over non-adnexal enriched (blue) samples. **J)** Myeloid cell cluster composition panels analogous to C). Ranked by fraction of dendritic cells (left), M1.S1008 cells (middle) and M2.SELENOP cells (right). \*\**P*<0.01, \*\*\**P*<0.001.

To identify the differentiation trajectories of the cell populations identified by unsupervised clustering, we employed diffusion manifolds to map CD8^+^ T cell clusters along two principal divergent branches representing the terminally differentiated populations, namely dysfunctional T cells and T cells expressing interferon-stimulated genes (ISG) (**Fig. 3D**). Assessment of scaled module scores with respect to differentiation pseudotime identified progressive loss of expression of naive T cell markers and progressive acquisition of cytotoxic and dysfunctional traits (**Fig. 3E**). These, in turn, were associated with progressive loss of expression of transcription factors associated with naive and central memory T cells (*TCF1* and *LEF1*), and gradual acquisition of gene expression related to type I IFN (*ISG15*), cytotoxic function (*GZMK*), and T cell dysfunction (*TOX*, *CXCL13*, and *PDCD1*) (**Fig. S3B**). Similar to the findings from T cell cluster abundance, expression gradients also differed across sites, with ascites samples exhibiting high naive module scores, contrasted by low dysfunctional T cell and JAK-STAT pathway scores (**Fig. 3F**).

### Macrophages and dendritic cells exhibit site-specific phenotypic enrichment

We next analyzed the composition and site distribution of the remaining major immune cell types. We identified four different dendritic cell (DC) states (**Fig. 3G**), including DCs of the myeloid lineage separated into cDC1, cDC2 and mature conventional dendritic cells (mDC), defined by expression of *CLEC9A, S100B and BIRC3* respectively (**Fig. S3C**, **Tab. S3**), and plasmacytoid DCs (pDC), marked by expression of *PTGDS*. Macrophage clusters were described with respect to their classical (M1-type) or alternative (M2-type) polarization. Six different clusters encompassing both classical and alternatively-activated macrophages were identified, as well as a cluster of cycling (Cycling.M) and a cluster of actively phagocytic macrophages (Clearing.M) (**Fig. 3G**). The M1-type and M2-type clusters were labeled according to the top genes defining the clusters (M1.S100A8, M2.CXCL10, M2.SELENOP, M2.MARCO, M2.COL1A1, M2.MMP9) (**Fig. S3C**, **Tab. S3**) ^20,21^. Notably, the M2.CXCL10 cluster was characterized by expression of both M1 (e.g. *CXCL10*) and M2 markers (e.g. PDL1, *C1QC*), highlighting that macrophage polarization represents a dynamic range of macrophage states rather than discrete functional phenotypes.

Both GLM and kernel density estimates highlighted inter-site differences in the relative composition of some of the largest myeloid cell clusters (2, 6, 8) (**Fig. 3H,I**) including DC enrichment in the ascites (log_2_ odds = +0.89), and depletion in the adnexa (**Fig. 3J** left, **Fig. S3D,** log_2_ odds = -0.21). Similarly, among macrophages, M1.S100A8 fractions were decreased in adnexa (log_2_ odds = -0.41) and increased in ascites samples (log_2_ odds = +1.50), while M2.SELENOP fractions demonstrated an opposite pattern (**Fig. 3J** middle and right), depleted in ascites (log_2_ odds = -1.28) and suggestive of a more immunosuppressive TME in the adnexa (log_2_ odds = +0.50). Altogether, our analyses of the T cell and myeloid cell compartments highlight that intra-patient variation in the tumor immune composition is common in HGSOC, with adnexal sites exhibiting evidence of reduced immune response and enrichment for specific functional immunosuppressive states.

### Mutational processes impact cancer cell intrinsic signaling states and expression of immuno-modulatory genes

We next focused on cancer cells to identify distinct cell-intrinsic phenotypic states. After normalization and regressing out patient-specific variation, clustering of 251,837 epithelial cells from scRNA-seq resulted in 10 clusters with cell-intrinsic differential expression of known marker genes (**Fig. 4A, Tab. S3**). Elevated TNFα, NF-κB and JAK-STAT signaling was observed in Cancer.cell.3 (including *CXCL10* and *ISG15*), elevated TGFβ signaling in Cancer.cell.4, and hypoxia in Cancer.cell.6. We note parenthetically that p53 signaling activity in Ciliated.cell.1 and Ciliated.cell.2 suggested presence of non-malignant epithelial cells in the tumor microenvironment. Distinct cluster enrichments were identified in association with patients stratified by genomic subtype (**Fig. 1B, Fig. 4C,D, Fig. S4A**): Cancer.cell.3 in HRD-Dup (log_2_ odds = +1.17), Cancer.cell.2 in HRD-Del (log_2_ odds = +0.34), and Cancer.cell.6 in FBI cases (log_2_ odds = +0.26). Among the three pathways with high activity in Cancer.cell.3, JAK-STAT signaling was significantly increased across HRD-Dup cases compared to HRD-Del and FBI cases (**Fig. 4E**, **Fig. S4B**, *P* = 0.0034; 0.026). NF-κB and TNFα signaling activity was increased in HRD-Dup and HRD-Del patients over FBI patients (*P* = 0.0073; 0.017; 0.012; 0.096) while TGFβ signaling was higher in FBI patients (*P* < 0.0001). In addition, cancer cell clusters differed by expression of major histocompatibility complex (MHC)-encoding genes (**Fig. 4F** left, **Fig. S4D,E**). MHC class I encoding genes (*HLA-A*, *HLA-B*, *HLA-C* and *B2M*) were highly expressed in Cancer.cell.3 (log_2_ fold changes of 0.77, 0.81, 0.69 and 0.78 respectively). MHC class II encoding genes (*HLA-DRA* and *HLA-DRB1*) were also increased (log_2_ fold changes of 0.36, 0.41). Consistent with higher Cancer.cell.3 distributions in HRD-Dup, we observed upregulation of *HLA-A*, *HLA-DRA*, *HLA-DRB1*, and *B2M* in HRD-Dup relative to FBI tumors (**Fig. 4F**, right). Whereas *HLA* gene expression suggests that Cancer.cell.3 cluster may be more immunogenic, we also noted upregulated expression of *CD274* which encodes PDL1 (**Fig. 4G**, *P* = 0.0028). Together, these findings imply that underlying genomic subtypes are associated with differential cancer cell-intrinsic signaling activation, with increased JAK-STAT signaling and upregulation of immune-activating and immune-inhibitory targets being more prevalent in HRD-Dup subtype.

**Figure 4.**
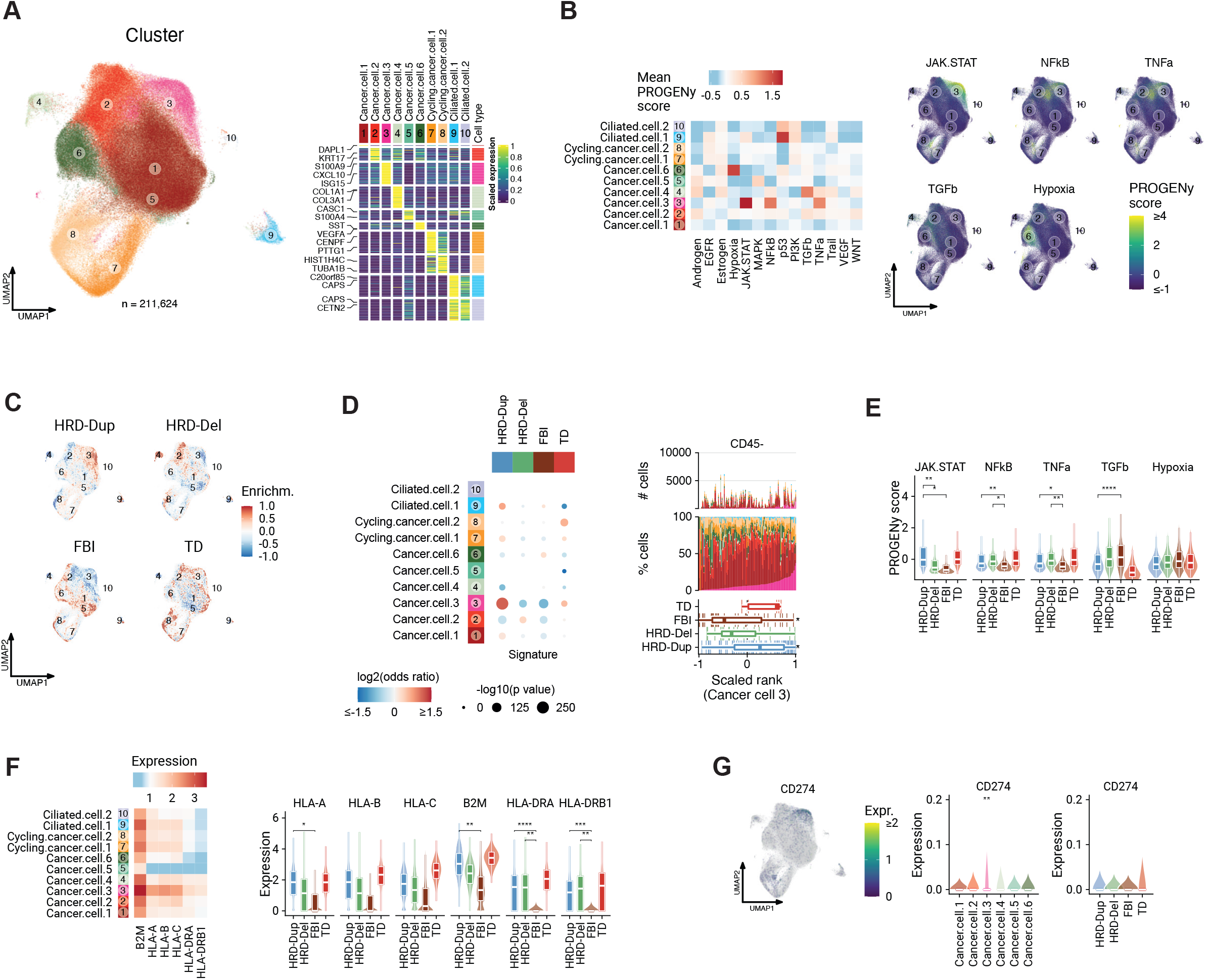
HR deficiency alters the landscape of cancer cell signaling states. **A**) Left: UMAP of epithelial cells colored by cluster. Clusters are numbered to reference cluster labels in heatmap. Right: Heatmap of scaled marker gene expression (averaged per cluster), showing differentially expressed genes in rows and clusters in columns. Top 2 genes per cluster are highlighted. **B)** Left: Heatmap of average signaling pathway activity scores. Right: UMAP colored by signaling pathway activity scores of interest. **C)** Relative kernel densities showing enrichment (red) and depletion (blue) in UMAP space for a given mutational signature. **D)** Left: Estimated effects of mutational signature on cancer cell cluster composition based on GLM. Color gradient indicates log_2_ odds ratios (enrichment: red, depletion: blue) and sizes indicate the BH-corrected -log_10_(p value). Right: Epithelial cluster compositions ranked by Cancer.cell.3 fractions. Box plot panels show distributions of scaled sample ranks by mutational signature. **E)** Distributions of signaling pathway activity scores with respect to mutational signature. **F)** Left: Heatmap of average HLA gene expression across clusters. Right: Distributions of HLA gene expression with respect to mutational signature. **G)** *CD274* (PDL1) gene expression in UMAP space (left) and as box plot distributions (right) with respect to cluster and mutational signature respectively. \**P*<0.05, \*\**P*<0.01, \*\*\**P*<0.001, \*\*\*\**P*<0.0001. Brackets: Wilcoxon pairwise comparisons.

### Genomic and microenvironment properties of HRD and FBI tumors identify mechanisms of immune recognition and escape

To evaluate the genomic and TME features conferring increased immunogenicity to HRD tumors, we evaluated the presence of novel mutant peptides, or neoantigens, in each genomic subtype. Using WGS, we predicted neoantigen burden on the basis of increased binding affinity of mutant peptides to patient-specific HLA types when compared to wild-type peptides, resulting in median 7 neoantigens per tumor sample (range 0 to 36, **Fig. 5A**). HRD-Dup and HRD-del cases exhibited significantly higher neoantigen burden (median 9 and 20.5 per sample, respectively) relative to FBI and TD cases (3 and 7 per sample, respectively, *P* < 0.05) (**Fig. 5B**), suggesting that inactivation of DNA repair triggers the accumulation of neoantigens. We then examined the relationship between genomic subtype and functional states of the tumor-infiltrating immune cells. Notably, mutational signatures were associated with relative proportions of naive and dysfunctional T cells through kernel density estimates in UMAP space (**Fig. 5C**). Quantitative modeling of T cell cluster compositions using GLM revealed FBI tumors were enriched for naive/stem-like and central memory T cell clusters (2-3, 15-16; log_2_ odds_FBI_ = +0.29) and depleted for dysfunctional T cell clusters (4-6, 18-20; log_2_ odds_FBI_ = -0.42) (**Fig. 5D**, **Fig. S5C**). Conversely, HRD and TD tumors were enriched for dysfunctional T cells (log_2_ odds_HRD-Dup_ = +0.17, log_2_ odds_HRD-Del_ = +0.39, log_2_ odds_TD_ = +0.15) and depleted for naive/stem-like and central memory T cells (log_2_ odds_HRD-Dup_ = -0.23, log_2_ odds_TD_ = - 0.23). This was also reflected in the enrichment of module scores across samples in general (**Fig. 5E**, **Fig. S5A**), and along differentiation trajectories of T-cell phenotypes in particular (**Fig. 5F**). In addition, the ISG-expressing T cell group (defined by high JAK-STAT pathway score) was negatively associated with FBI and HRD-Del subtypes, but was positively associated with HRD-Dup and TD subtypes (**Fig. 5E,F, Fig. S5B**). Comparison of JAK-STAT signaling in CD8^+^ T cells and matched cancer cells from the same samples, revealed strong association between T cell-intrinsic and cancer-cell intrinsic JAK-STAT signaling across the mutational subtypes (**Fig. 5G**).

**Figure 5.**
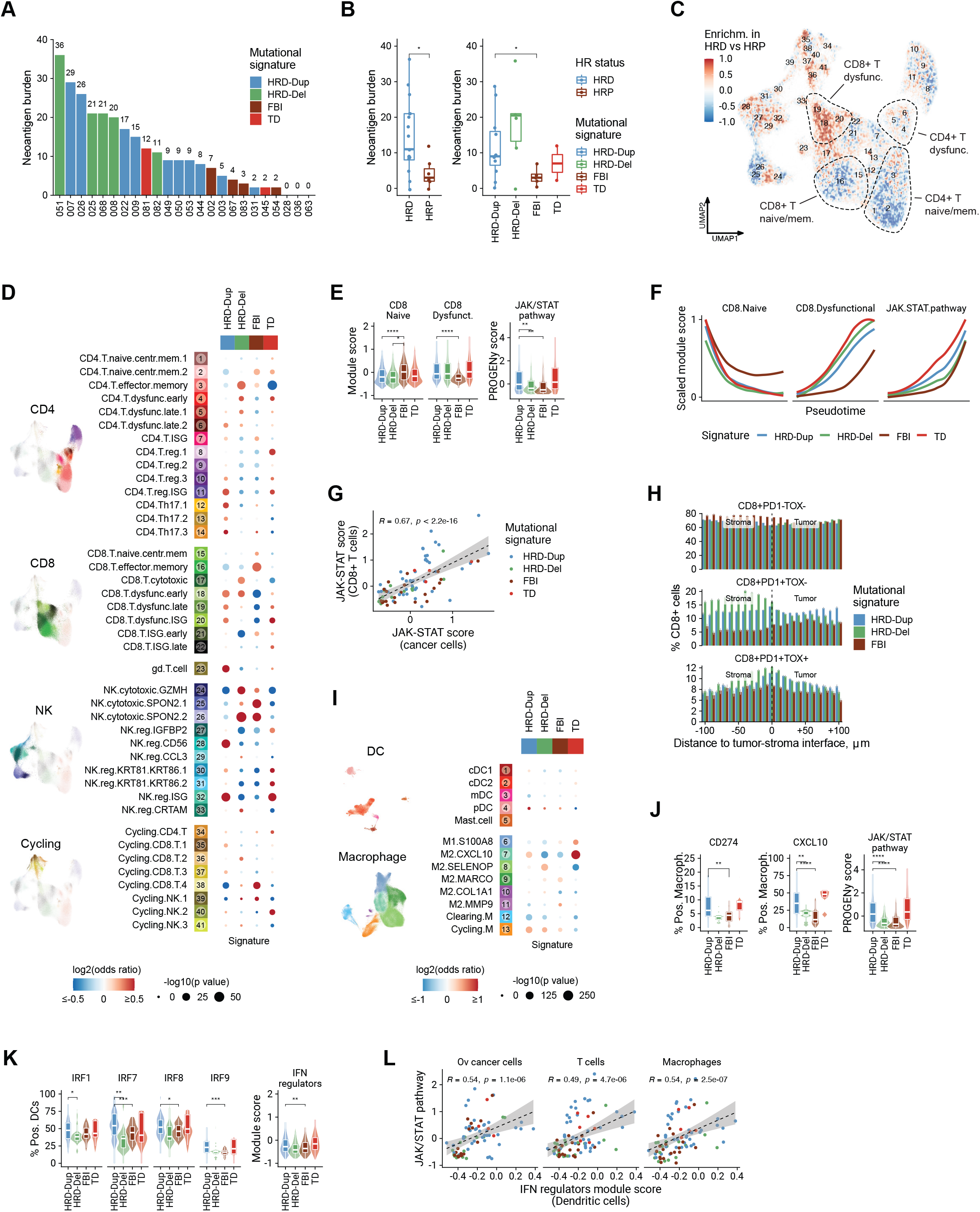
HR deficiency is associated with increased antigen burden and dysfunctional T cell phenotypes. **A)** Neoantigen burden detected by whole genome sequencing. Bar graphs indicate the total count per tumor sample of neoepitopes exhibiting mutant binding affinity of <1,000 nM. **B)** Pairwise comparisons of neoantigen burden with respect to HR status and mutational signature. **C)** Differences in kernel density estimates in UMAP space for HRD (red) over HRP (blue) samples. **D)** Estimated effects of mutational signature on T and NK cell cluster composition based on GLM. Color gradient indicates log_2_ odds ratios (enrichment: red, depletion: blue) and sizes indicate the BH-corrected -log_10_(p value). **E)** Distributions of CD8^+^ T cell state module scores and JAK-STAT signaling pathway activity scores with respect to mutational signature. **F)** Scaled module scores within the subset of CD8^+^ T cells with respect to pseudotime and mutational signature. **G)** Correlation of JAK-STAT signaling scores in CD8^+^ T cells in CD45^+^ samples and scores in cancer cells in matched CD45^-^ samples. **H)** Spatial density of CD8^+^ T cell phenotypes as a function of distance to the tumor-stroma interface, grouped by mutational signature (**Methods**). **I)** Estimated effects of mutational signature on myeloid cell cluster composition using GLM, analogous to D). **J)** Fraction of macrophages expressing *CD274* (PDL1) and *CXCL10* and JAK-STAT pathway activity with respect to mutational signature. **K)** Left: Fraction of dendritic cells expressing interferon regulating factors. Right: Module score of all IFN regulators. **L)** Correlation of IFN regulators expressed in dendritic cells with JAK-STAT signaling scores in cancer cells, T cells and macrophages. \**P*<0.05, \*\**P*<0.01, \*\*\**P*<0.001, \*\*\*\**P*<0.0001. Brackets: Wilcoxon pairwise comparisons.

We next tested if heightened immune signaling in HRD tumors could be attributed to reciprocal interactions between cancer cells and the associated immunophenotypes in the TME. We analyzed the physical proximity of individual T cell phenotypes to the tumor-stroma boundary using the *in situ* mpIF data obtained from the site-matched tumor samples. Cell density estimates inferred from site-matched mpIF data as a function of distance to the tumor-stroma boundary revealed that CD8^+^ T cells were enriched within tumor regions of HRD subtypes (**Fig. S5D**), among which activated CD8^+^PD1^+^TOX^-^ T cells were especially prevalent (**Fig. 5H**). Similarly, terminally dysfunctional CD8^+^PD1^+^TOX^+^ T cells were also co-localized within the tumor in HRD-Dup and HRD-Del cases, and also in the peritumoral stroma of HRD-Del cases, potentially reflecting tumor reactivity. In contrast, both CD8^+^PD1^+^TOX^-^ and CD8^+^PD1^+^TOX^+^ T cells were less abundant and were evenly distributed within the tumor and stromal compartments in the FBI tumors, implying minimal T cell-antigen interaction in this subtype (**Fig. 5H**).

The balance of M1- and M2-type macrophage polarization states was similarly shaped by tumor mutational signatures (**Fig. 5I, Fig. S5E**), with a significant depletion of M2 macrophages in HRD-Del cases (log_2_ odds_HRD-Del_ = -0.27). We observed enrichment of M2.CXCL10 macrophages in the HRD-Dup cases (log_2_ odds_HRD-Dup_ = +0.33), and depletion in the FBI tumors (log_2_ odds_FBI_ = -0.46), with FBI tumors also exhibiting decreased percentages of PDL1 (*CD274*) positive macrophages (**Fig. 5J**). Consistent with *CXCL10* being a target of JAK-STAT signaling, macrophages in HRD-Dup cases were enriched for JAK-STAT pathway activity, whereas most FBI cases were depleted for JAK-STAT activation (**Fig. 5J**).

Overall, these findings imply that JAK-STAT pathway activation in all cell subtypes in the HRD-Dup tumors may be mediated by a common upstream effector. Given the role of type I IFNs in activation of JAK-STAT signaling, we examined type I IFN pathway activation in DCs, which commonly serve as a dominant source of type I IFN production. We observed higher levels of IFN regulatory factors (IRFs) in DCs from HRD-Dup cases, resulting in upregulation of the overall IFN regulators module score (**Fig. 5K**). There was a strong positive correlation between the IFN regulators module score in the DCs and JAK-STAT pathway activation in cancer cells, T cells, and macrophages (**Fig. 5L**). These findings may suggest that increased type I IFN activation in DCs in HRD-Dup tumors may be responsible for the upregulated JAK-STAT signaling and phenotypic changes in all major cell types, including HLA upregulation in cancer cells and PDL1 upregulation in macrophages.

### Immunoediting in HR-deficient tumors is mediated by HLA loss of heterozygosity

We next determined if genomic mechanisms of immune evasion across the mutational signature subtypes could explain the variation in immunophenotypes. Genomic loss of HLA presentation machinery through somatic alterations is a common mechanism for malignant cells to evade immune control by reduction of MHC class I allelic diversity ^2^. To examine how somatic HLA diversity varies as a function of genomic subtype, we identified allele-specific copy number alterations from scRNA-seq using heterozygous SNPs from matched normal whole genomes (**Methods**). Aggregating counts of heterozygous SNPs across each chromosome arm we inferred loss of heterozygosity (LOH) of either allele of chromosome arm 6p, harboring HLA class I and class II genes, at the single cell level (**Fig. 6A**). As expected, LOH of the 6p arm were only detected in cancer cells, indicating high specificity of the method (**Fig. 6A,B**, **Fig. S6A**). B-allele frequency (BAF) estimates revealed marked inter-patient heterogeneity in 6p allelic imbalance (**Fig. 6B**, **Fig. S6B**). Clonal LOH of chromosome 6p was seen in 4 out of 41 patients (10%), and subclonal 6p LOH was observed in 7 out of 41 patients (17%, **Fig. 6C** left). The prevalence of site-specific 6p LOH (**Fig. 6C** right) was low within most patients, with the exception of site-specific losses in 4 out of 41 cases (**Fig. 6C** right, **Fig. S6B**) indicating potential selection for immune evasive mechanisms determined by local microenvironments. We note that clonal LOH of 6p was primarily observed in HRD cases (17%, 4/24 cases) (**Fig. 6C,D**). Losses of HLA class I alleles harbored in 6p were validated using clinical panel sequencing of the MSK-IMPACT HGSOC cohort (n=1,111 patients). Patients with *BRCA1*, *BRCA2*, and *CDK12* loss of function mutations and *CCNE1* amplification were identified as representative cases of HRD-Dup, HRD-Del, TD and FBI signature groups, respectively. Overall, we observed HLA LOH in 333 out of 1,111 MSK-IMPACT HGSOCs (30%) (**Fig. 6D**, **Fig. S6C**). *BRCA1*-altered HGSOC cases showed more frequent HLA LOH than *CCNE1*-amplified cases (30% vs 22% of altered cases; *P* < 10^−13^, two-sided *χ*^2^ test), consistent with the findings identified by scRNA-seq. Validation of 6p BAF estimates from scRNA-seq were additionally corroborated by concordant HLA LOH status in site-matched MSK-IMPACT tumor samples (**Fig. S6D**).

**Figure 6.**
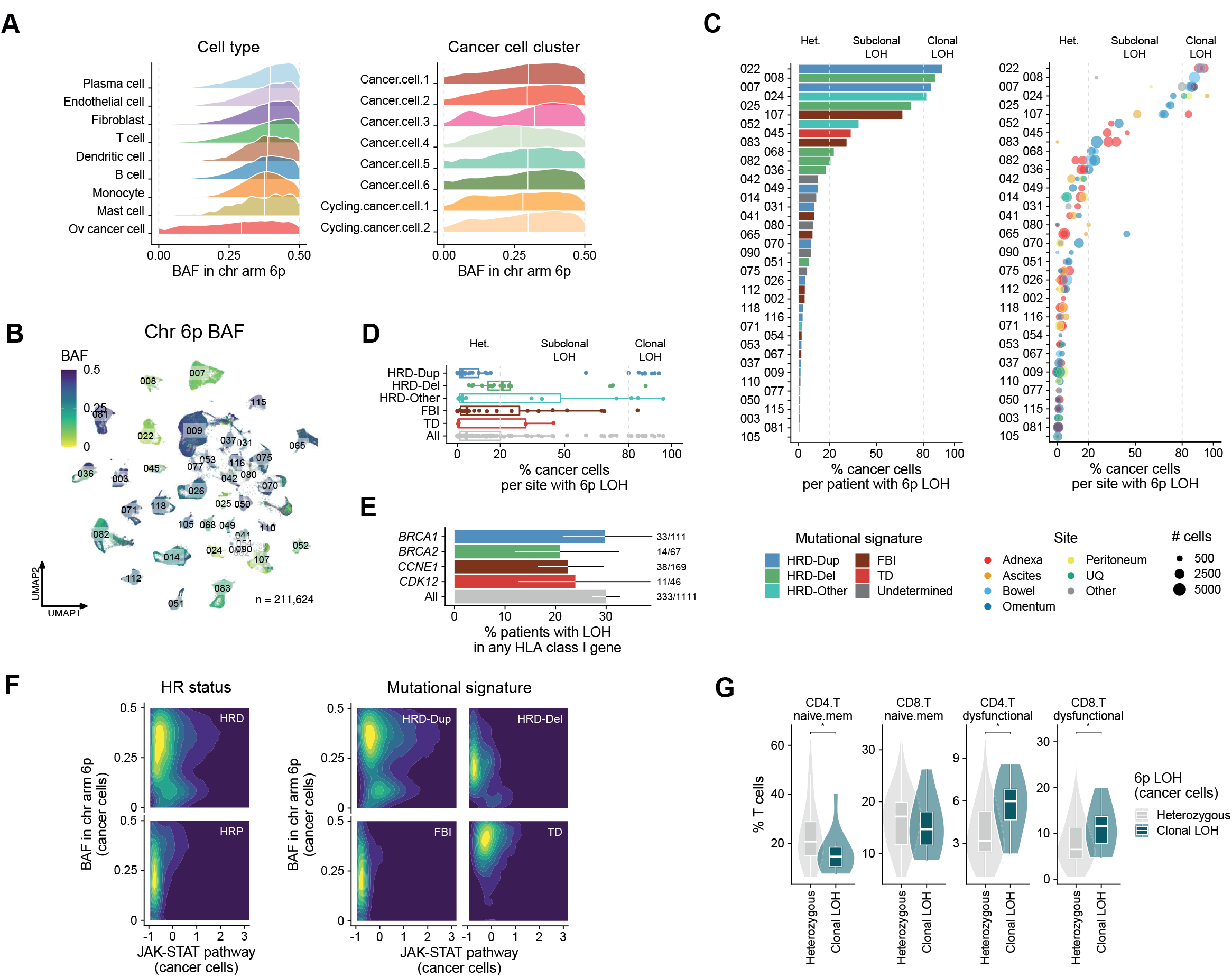
HR deficiency is associated with tumor immunogenicity and immune evasion. **A)** Density distribution of 6p BAF per cell in cancer cells compared to non-malignant cells, ranked by median 6p BAF per cell type (left panel). Allelic imbalance in 6p BAF across cancer cell clusters (right panel). White vertical lines indicate the median 6p BAF. **B)** UMAP of cancer cells profiled by scRNA-seq colored by BAF of chromosome arm 6p. **C)** Left: Percentage of cancer cells with LOH in chromosome 6p per patient. Right: Site- and clone-specific percentage of 6p LOH in cancer cells. **D)** Distributions over percentage of 6p LOH in cancer cells per sample as a function of mutational signature subtype. **E)** Percentage of patients with HLA LOH of any HLA class I gene in the MSK-IMPACT HGSOC cohort (n=1,111 patients) for *BRCA1*, *BRCA2, CDK12* mutant and *CCNE1* amplified tumors, mapping to HRD-Dup, HRD-Del, TD, and FBI signatures respectively. Error bars are 95% binomial confidence intervals **F)** Normalized density contours of 6p BAF and JAK-STAT pathway activity in cancer cells comparing HR status and mutational signature. **G)** Fraction of naive and dysfunctional T cells as a function of clonality of 6p LOH in cancer cells. Percentage allelic loss of chromosome 6p arm in cancer cells per site is used to bin samples according to their 6p LOH status (heterozygous: % 6p LOH < 20%, clonal LOH: % 6p LOH > 80%). In panels A)-F), only BAF estimates from cells with ≥10 reads aligning to chromosome arm 6p are considered, and allelic imbalance states are assigned per cell based on the mean 6p BAF per cell as balanced (BAF ≥ 0.35), imbalanced (0.15 ≤ BAF < 0.35) or LOH (BAF < 0.15) (**Methods**). \**P*<0.05. Brackets: Wilcoxon pairwise comparisons.

We then analyzed whether 6p LOH impacted tumor cell-intrinsic JAK-STAT pathway activation and phenotypic states of tumor-infiltrating T cells. We found that HRD-Dup tumors with 6p LOH frequently exhibited concomitant activation of JAK-STAT signaling (**Fig. 6E**, **Fig. S6E**). Furthermore, presence of dysfunctional CD4^+^ and CD8^+^ T cells was increased in tumors with clonal 6p LOH (**Fig. 6F**), consistent with loss of allelic diversity at the chromosome 6p locus resulting from evolutionary selective pressure exerted by infiltrating T cells. In contrast, the microenvironment of tumors with high prevalence of naive T cells did not impose a strong selective pressure of immune predation, evidenced by the retention of both 6p alleles in cancer cells (**Fig. 6F**). M1 macrophages and DCs were similarly more frequently observed in tumors with 6p LOH, though the association was not statistically significant (**Fig. S6F**).

### Spatial topology and site composition influence malignant cell selection and immune pruning

The single-cell analyses above point to an interrelationship between the cancer cell-intrinsic immune signaling, TME composition, and phenotypes of tumor-infiltrating immune cells that vary as a function of the underlying mutational signatures. Accordingly, cell-cell interaction analysis revealed co-expression of PDL1 (*CD274*) in cancer cell and myeloid clusters detected by scRNA-seq, and PD1 (*PDCD1*) in T and NK cell clusters (**Fig. 7A**). We noted higher expression of PDL1 in myeloid cell clusters and in Cancer.cell.3 cluster in HRD-Dup cases, associated with higher fraction of PD1-expressing T cells (**Fig. 7A**). Using mpIF stains from site-matched FFPE sections highlighting the principal immune cell subtypes and their phenotypic states, we then investigated whether these interrelationships are reflected in the *in situ* interactions between the individual cell components in tumors and stroma. We measured major immune cell subtypes (tumor cells, T cells, macrophages), markers of T cell dysfunctional states (PD1, TOX), and PDL1 expression as a functional downstream marker of type I and type II IFN signaling (**Fig. 7B**).

**Figure 7.**
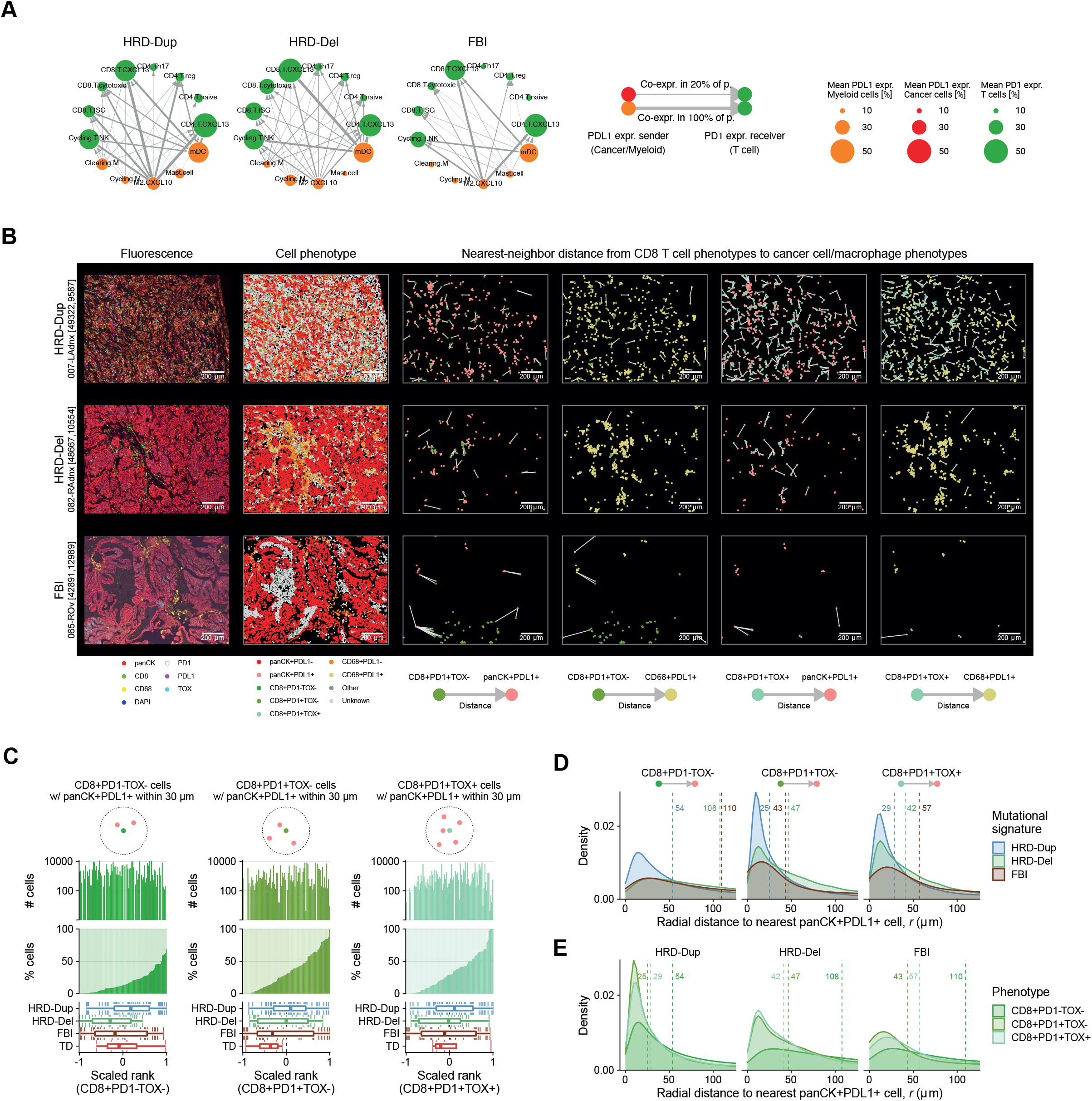
Spatial topology and site composition influences malignant cell selection and immune pruning. **A)** Interaction network diagrams depicting ligand-receptor co-expression across cell types faceted by mutational signature. Nodes show mean PD1 (*PDCD1*) expression in CD4^+^ T, CD8^+^ T and NK clusters, and mean PDL1 (*CD274*) expression in cancer cell and myeloid cell clusters in scRNA data, depicted by circle size. Arrows join ligand-expressing sender clusters to receptor-expressing receiver clusters and are weighted by frequency of co-expression of PD1 and PDL1 in sender and receiver clusters. **B)** Representative mpIF fields of view highlighting common features of the tumor microenvironment of mutational signature subtypes. First column: Raw pseudocolor images; second column: cellular phenotypes of segmented cells; remaining columns: proximity of pairs of phenotypes, highlighting ligand-receptor interactions between PDL1 and PD1 with color-coded phenotypes, and edges depicting distances. Only edges joining pairs of cells within 250 μm are shown. **C)** Proximity analysis between CD8^+^ T cell phenotypes and panCK^+^PDL1^+^ cancer cells based on mpIF data, ranking samples by the fraction of CD8^+^PD1^-^TOX^-^ T cells (left), CD8^+^PD1^+^TOX^-^ T cells (middle) or CD8^+^PD1^+^TOX^+^ T cells (right) with ≥1 panCK^+^PDL1^+^ cell within 30 μm. Upper panels: absolute abundance of CD8^+^ T cell states; middle panels: fraction of CD8^+^ T cell phenotypes with ≥1 panCK^+^PDL1^+^ cell within 30 μm; bottom panels: box plot distributions of sample ranks with respect to mutational signature. **D and E)** Nearest-neighbor distance from CD8^+^ T cell phenotypes to panCK^+^PDL1^+^ cancer cells aggregated across fields of view, grouped by mutational signature subtype. Vertical lines indicate the median nearest-neighbor distance.

Nearest-neighbor analysis between cells revealed proximal interaction patterns between CD8^+^PD1^-^ TOX^-^, CD8^+^PD1^+^TOX^-^, CD8^+^PD1^+^TOX^+^ T cells, defined as naive/memory, activated/pre-dysfunctional and dysfunctional T cells, respectively, and PDL1-expressing cancer cells and macrophages (panCK^+^PDL1^+^, CD68^+^PDL1^+^) within a 30 μm radius (**Fig. 7C**, **Fig. S7A**). Spatial proximity of antigen-experienced PD1^+^ T cells to PDL1^+^ cancer cells was commonly observed in HRD-Dup and HRD-Del cases. However, these interactions were rare or absent in FBI tumors. The overall spatial organization, median distances between the individual T cell subtypes and the nearest panCK^+^PDL1^+^ cancer cells varied as a function of the mutational type, with the closest median distances observed in the HRD-Dup subtype, particularly in the activated/pre-dysfunctional and dysfunctional T cell compartments (**Fig. 7D,E**). The proximity of pre-dysfunctional and dysfunctional T cells to PDL1-expressing cancer cells supports the hypothesis that PDL1 is a functional biomarker induced as a negative feedback mechanism in response to activated T cells in HRD tumors. Similar interactions were noted between the T cells and CD68^+^PDL1^+^ macrophages, which were particularly prevalent in the HRD-Del cases, likely due to their overall enrichment for macrophages (**Fig. S7B,C**, **Fig. 5H**). These spatial associations in HRD subtypes revealed a highly immunoreactive TME, where PDL1 induction in cancer cells and macrophages was prevalent within ∼50 μm of T cells (**Fig. 7E**). Conversely, spatial correlations between T cells and PDL1^+^ cancer cells quickly decayed in FBI cases, reflecting insufficient T cell activation to mediate the local TME changes.

## DISCUSSION

Here, we outline both anatomic site and mutational processes as determinants of within and between patient variation of immune cell functional states in HGSOC. Our study reveals for the first time distinct immune evasion phenotypes in HR-deficient and HR-proficient HGSOC genomic subtypes. The marked degree of intra-patient tumor microenvironment heterogeneity across tumor sites highlights that interfaces of cancer cells and their surrounding immune compartment can be site-specific within patients. Specifically, adnexal sites exhibit the lowest levels of T cell infiltration and a marked increase in T cell dysfunction. The biology underlying distinct immunophenotypes in adnexal samples is unclear. However, our results may indicate immune privilege of the ovaries and fallopian tubes as protective environments against inflammation and immune pruning. Furthermore, we show that cancer and immune cell states in the ascites samples are distinct from the rest of the tumors, implying that the phenotypes of ascitic sample cells might not reflect intact tumor tissues. We find that ascites are enriched for T cells exhibiting predominantly naive-like and central memory signatures, while being depleted for activated and dysfunctional T cells. This is supported by prior findings^22^ and is consistent with the requirement for chronic antigen stimulation to elicit differentiation of dysfunctional T cells. Similarly, we find that ascitic fluid is uniquely enriched for cDC1 and cDC2 dendritic cell subsets relative to tumors, consistent with recent findings demonstrating that ascitic myeloid cells can be rapidly adapted to serve as antigen-presenting cells in vaccine preparation^23^.

Most notably, our study demonstrates that different mutational processes associated with HGSOC engender distinct immune evasion mechanisms. FBI tumors exhibit overall low immunogenicity associated with low neoantigen burden, reduced expression of HLA genes, and increased TGFβ signaling activity, with predominance of naive/stem-like and central memory T cells, likely driven by exclusion of T cells from the tumor compartment. In contrast, HRD tumors demonstrate evidence of intrinsic immunogenicity, including neoantigen accrual, upregulation of JAK-STAT/type I IFN signaling and increased expression of HLA genes, with associated infiltration of tumors with T cells exhibiting signatures of dysfunction, likely elicited by chronic interaction with tumor antigens. Critically, as FBI tumors have worse prognosis, these tumors represent a high-risk group that are immunologically inert and thus should be investigated as a high priority group of patients for emerging immuno-stimulatory agents.

Data from preclinical studies of ovarian cancer suggests that immunogenicity in HRD tumors may lead to improved responses to immune checkpoint blockade (ICB) or combinations of ICB with chemotherapy and PARP inhibitors ^24–27^. However, clinical evidence for this is lacking. In our recent analysis of ovarian cancer patients treated with ICB, no association between HRD status, tumor mutational burden (TMB) and response to immunotherapy was observed ^28^. Furthermore, two large randomized clinical trials evaluating combination of platinum-based chemotherapy with PDL1 inhibition in the newly-diagnosed ovarian cancer patients failed to identify benefit from the addition of PDL1 inhibitor ^29,30^, while subset analyses from at least one of the trials showed no evidence of improved response in the subset of patients carrying tumors with evidence of *BRCA1/2* mutations (Lederman et al., IGCS 2020). These findings highlight that data from pre-clinical models may not accurately reflect the evolution of human HGSOCs, where despite evidence of apparent immunogenicity, HR-deficient tumors potentially develop antagonizing mechanisms of immune resistance. Furthermore, our data are consistent with distinct immune escape mechanisms between non-HRD and HRD tumors, thus indicating that distinct strategies for immunotherapeutic reprogramming may be required for these cancer types. Increased TGFβ signaling observed in FBI tumors has been shown to restrict T-cell infiltration and is associated with insensitivity to anti-PDL1 treatment ^31,32^. Hence, blocking TGFβ signaling in HGSOC patients of FBI subtype might help to overcome the immune exclusion, as has been suggested for TGFβ-high colorectal cancer patients^33^. In HRD tumors, the tumors exhibit inherent immunogenicity, consistent with ongoing immune response eliciting co-evolution of cancer cells with their microenvironment, and leading to immune evasion through selection of HLA LOH in these cancers. Lastly, a high degree of inter-tumor microenvironment heterogeneity in all genetic subtypes highlights that the mechanisms of immune resistance might not be universal in a given patient and that evolution of immune response in individual tumors may lead to acquisition of additional resistant mechanisms in these tumors, as evidenced by heterogeneity in HLA LOH observed across tumors in some patients and intrinsic differences in anatomic sites. Notably, tumor heterogeneity defined by radiomic measures has been demonstrated to be associated with resistance to immunotherapy in HGSOC ^34^.

In addition to well-studied mechanisms of immune evasion, we identified concomitant and potentially chronic upregulation of type I IFN signaling in T cells, cancer cells and cancer-associated myeloid cells, particularly enriched for tumors of HRD-Dup subtype. Expression of type I IFN signaling target genes in such a wide range of cell types implies a potentially common source of type I interferons eliciting these signatures in all cell populations. The significance of this finding at present is unknown, but possibly related to cancer cell-intrinsic genomic instability eliciting type I IFN production through generation of cytosolic DNA or upregulation of noncoding elements ^35,36^. As chronic type I IFN production is associated with upregulation of immune suppressive mechanisms and T cell dysfunction, particularly within the context of chronic virus infections and cancers ^37,38^, we suspect that this finding represents yet another mechanism driving the differentiation of dysfunctional T cells and immune escape.

Altogether, our study provides an extensive multi-modal resource mapping the cellular constituents of HGSOC tumor microenvironments and linking them to mutational processes and spatial context. We suggest these findings represent an opportunity to understand the adaptive immune response in other cancers with genomic instability through similar approaches. Whether anatomic site of tumor foci or structural variation mutational processes in other cancers are determinants of immune response remain open questions. However, the data presented here can be leveraged to contextualize mechanistic insights of immunotherapeutic response across other cancers of genomic instability.

### Data availability

Raw sequencing data and gene expression counts for 10x 3’ scRNA-seq will be available from the NCBI GEO database prior to publication. MSK-IMPACT and WGS data will be released on the NCBI SRA and on cBioPortal upon publication. H&E and mpIF images will be available from the BioImage Archive.

## Acknowledgements

This project was funded in part by Cycle for Survival supporting Memorial Sloan Kettering Cancer Center. SPS holds the Nicholls Biondi Chair in Computational Oncology and is a Susan G. Komen Scholar. This work was funded in part by awards from the Ovarian Cancer Research Alliance (OCRA) Collaborative Research Development Grant to SPS [648007], OCRA Liz Tilberis Award to DZ [657721] and OCRA Ann Schreiber Mentored Investigator Award to IVG [650687], the Department of Defense Congressionally Directed Medical Research Programs W81XWH-20-1-0565 to SPS, DZ and BW, the Department of Defense Ovarian Cancer Research Academy OC150111 to DZ, the LesLois Shaw Foundation and the Cancer Research UK Grand Challenge Program to SPS [C42358/A27460], the Marie-Josée and Henry R. Kravis Center for Molecular Oncology and the National Cancer Institute (NCI) Cancer Center Core Grant No. P30-CA008748. KMB is supported by NCI F30 CA257414 and the NIH T32 MD-PhD training program GM007739. FP is supported by NCI K12 CA184746.

## Declaration of interests

SPS is a consultant and shareholder of Canexia Health Inc. DZ reports research funding to MSK from AstraZeneca, Genentech, and Plexxikon; personal fees from Synlogic Therapeutics, Hookipa Biotech, Agenus, Synthekine, Memgen, Mana Therapeutics, Tessa Therapeutics, and Xencor; stock options from Accurius, Calidi Biotherapeutics, and Immunos. DZ is an inventor on a patent concerning the use of Newcastle Disease Virus as a cancer therapeutic, licensed to Merck. CFF reports research funding to the institution from Merck, AstraZeneca, Genentech, Bristol Myer Squibb, Taiho and Seattle Genetics; uncompensated membership of a scientific advisory board for Merck and Genentech; and is a consultant for OncLive, Aptitude Health, and Seattle Genetics, all outside the scope of this manuscript. CA reports grants from Clovis, Genentech, AbbVie and AstraZeneca, personal fees from Tesaro, Eisai/Merck, Mersana Therapeutics, Roche/Genentech, Abbvie, AstraZeneca/Merck and Repare Therapeutics, outside the submitted work. NRAR reports grants to MSK from Stryker/Novadaq and GRAIL, outside the submitted work. DSC is on the medical advisory board of Apyx Medical Co, Verthermia Acquio Inc and Biom’up, and is a stockholder of Intuitive Surgical Inc and TransEnterix Inc. RNG reports funding from GSK, Mateon Therapeutics, Regeneron, Clovis, Context Therapeutics, EMD Serono, MCM Education, OncLive, Aptitude Health and Prime Oncology, outside this work. YLL reports research funding from AstraZeneca and GSK/Tesaro outside this work. YL is a shareholder of Y-mAbs Therapeutics Inc. TJH receives research funding from Bristol Myers Squibb, Calico Labs and the Parker Institute for Cancer Immunotherapy. SFB owns equity in, receives compensation from, and serves as a consultant and the Scientific Advisory Board and Board of Directors of Volastra Therapeutics Inc. SFB has also consulted for Sanofi, received sponsored travel from the Prostate Cancer Foundation, and both travel and compensation from Cancer Research UK. BW reports ad hoc membership of the scientific advisory board of Repare Therapeutics, outside the submitted work.

## Author contributions

SPS, DZ: project conception and oversight; IVG, FU, NR, DZ, SPS: manuscript writing and editing; IVG, FU, SPS, DZ: study design; NAR, CA, CF, BW, MW, RN, JL: clinical research coordination; JL, MW, ADCP, AMaroldi, AStylianou, FND, FP, MV, BW: tissue procurement, biological substrates, data generation; DZ, YB, SHK: dissociation protocols; NM, AV: single cell RNA sequencing, genome sequencing; AR, TJH: multiplexed imaging; IVG, FU, NC, KMB, TF, AMcPherson, HS, MJW, AWZ: computational biology, data analysis; IVG, FU, MP, RV, DP, CF, LG: data curation; AMcPherson, DG, SL, EH, DP, AP, VBojilova, JG, RK, DK: data processing, visualization; NRAR, DSC, KLR, OZ, YS, GJG, VBroach: surgery; DZ, CA, CFF, YLL, ML, RNG: clinical data review; LHE, RAS: pathology review; YL, IN: radiology review; SFB, ASchietinger: discussion. All authors read and approved the final manuscript.

## METHODS

### Experimental methods

#### Sample collection

All enrolled patients were consented to an institutional biospecimen banking protocol and MSK-IMPACT ^39^, and all analyses were performed per a biospecimen research protocol. All protocols were approved by the Institutional Review Board (IRB) of Memorial Sloan Kettering Cancer Center. Patients were consented following the IRB-approved standard operating procedures for informed consent. Written informed consent was obtained from all patients before conducting any study-related procedures. The study was conducted in accordance with the Declaration of Helsinki and the Good Clinical Practice guidelines (GCP).

We collected fresh tumor tissues from 42 HGSOC patients at the time of upfront diagnostic laparoscopic or debulking surgery. Ascites and tumor tissue from multiple metastatic sites, including bilateral adnexa, omentum, pelvic peritoneum, bilateral upper quadrants, and bowel were procured in a predetermined, systemic fashion (median of 4 primary and metastatic tissues per patient) and were placed in cold RPMI for immediate processing. Blood samples were collected pre-surgery for the isolation of peripheral blood mononucleated cells (PBMCs) for normal whole genome sequencing (WGS). The isolated cells were frozen and stored at -80°C. In addition, tissue was snap frozen for bulk DNA extraction and tumor WGS. Tissue was also subjected to formalin fixation and paraffin-embedding (FFPE) for histologic, immunohistochemical and multiplex immunophenotypic characterization.

#### Single cell RNA sequencing

##### Tissue dissociation

Tumor tissue was immediately processed for tissue dissociation. Fresh tissue was cut into 1 mm pieces and dissociated at 37°C using the Human Tumor Dissociation Kit (Miltenyi Biotec) on a gentleMACS Octo Dissociator. After dissociation, single cell suspensions were filtered and washed with Ammonium-Chloride-Potassium (ACK) Lysing Buffer. Cells were stained with Trypan Blue and cell counts and viability were assessed using the Countess II Automated Cell Counter (ThermoFisher) (for detailed protocol see ^40^).

##### Cell sorting

Freshly dissociated cells were stained with a mixture of GhostRed780 live/dead marker (TonBo Biosciences) and Human TruStain FcX™ Fc Receptor Blocking Solution (BioLegend). The stained samples were then incubated and stained with Alexa Fluor® 700 anti-human CD45 Antibody (BioLegend). Post staining, they were washed and resuspended in RPMI + 2% FCS and submitted for cell sorting. The cells were sorted into CD45 positive and negative fractions by fluorescence assisted cell sorting (FACS) on a BD FACSAria™ III flow cytometer (BD Biosciences). Positive and negative controls were prepared and used to set up compensations on the flow cytometer. Cells were sorted into tubes containing RPMI + 2% FCS for sequencing.

##### Library preparation

Flow sorted tumor cells were stained with Trypan blue and Countess II Automated Cell Counter (ThermoFisher) was used to assess both cell number and viability. Following QC, the single cell suspension was loaded onto Chromium Chip B (10x Genomics PN 2000060). GEM generation, cDNA synthesis, cDNA amplification, and library preparation of 1,400-5,000 cells proceeded using the Chromium Single Cell 3’ Reagent Kit v3 (10x Genomics PN 1000075) according to the manufacturer’s protocol. cDNA amplification included 12 cycles and 0.4-419 ng of the material was used to prepare sequencing libraries with 8-14 cycles of PCR.

##### Sequencing

Equimolar amounts of indexed libraries were pooled and sequenced on a HiSeq 2500 in Rapid Mode or NovaSeq 6000 in a 28bp/91bp, 100bp/100bp, or 150bp/150bp paired end run using the HiSeq Rapid SBS Kit v2 or NovaSeq 6000 SP/S1/S2/S4 Reagent Kit (100/200/300 cycles) (Illumina).

#### Bulk whole genome sequencing (WGS)

##### WGS Bulk Tumor

Frozen banked tissue was cut into sections on charged microscope slides. Following histologic review, tumor tissue was microdissected if required to enrich for neoplastic cells ^41^, and subjected to DNA extraction for bulk WGS. Genomic DNA was extracted using the DNeasy Blood & Tissue kits (Qiagen) and quantified on the Qubit 3 Fluorometer using Qubit™ 1X dsDNA HS Assay Kit (Invitrogen).

##### WGS Bulk Normal

PBMCs were brought up to 15 mL volume in cold PBS and DNA was isolated with the DNeasy Blood & Tissue Kit (Qiagen catalog # 69504) according to the manufacturer’s protocol with 1 hour of incubation at 55°C for digestion. DNA was eluted in 0.5x Buffer AE.

##### Whole Genome Sequencing

DNA quantity was measured using the Quant-iT PicoGreen dsDNA Assay (ThermoFisher catalog # P11496), and DNA quality was assessed by TapeStation D1000 ScreenTape (Agilent catalog # 5067-5582). After PicoGreen quantification and quality control by Agilent BioAnalyzer, 500ng of genomic DNA were sheared using a LE220-plus Focused-ultrasonicator (Covaris catalog # 500569) and sequencing libraries were prepared using the KAPA Hyper Prep Kit (Kapa Biosystems KK8504) with modifications. Briefly, libraries were subjected to a 0.5x size select using aMPure XP beads (Beckman Coulter catalog # A63882) after post-ligation cleanup. Libraries were not amplified by PCR and were pooled equivolume and quantitated based on their initial sequencing performance. Samples were run on a NovaSeq 6000 in a 150bp/150bp paired end run, using the NovaSeq 6000 SBS v1 Kit and an S1, S2, or S4 flow cell (Illumina).

#### Preparation, review and scanning of histopathology slides

Archived formalin-fixed paraffin-embedded (FFPE) tissues were used for histologic review, including the assessment of spatial topology and tumor-infiltrating lymphocytes, as well as for immunohistochemical characterization and multiplex immunofluorescence analysis for mapping of the tumor microenvironment in the Advanced Immunomorphology Platforms Laboratory. Slides were originally reviewed by gynecologic pathologists for diagnosis and FIGO (International Federation of Gynecology and Obstetrics) stage assignment. Representative hematoxylin and eosin (H&E)-stained slides from each site of interest were digitally scanned to produce virtual slides. Two senior gynecologic pathologists (R.A.S., L.H.E.) then reviewed these images for the presence and location of serous tubal intraepithelial carcinoma (STIC), SET architecture (solid, pseudo-endometrioid and transitional cell-like patterns), micropapillary architecture ^42^, presence of a fimbrial ball, architectural patterns of metastatic disease ^43^, mitotic counts (per 10 high power fields, HPFs) and tumor cell content (viable %). Regions with tumor-infiltrating lymphocytes (TILs) were also assessed with a quantitative TIL score (low: <42 TILs per 1 HPF in a hotspot; high: 42 or more TILs per 1 HPF in a hotspot) ^42^. Histopathology slides were scanned into whole-slide images using a Leica Aperio AT2 scanner (Leica Biosystems) at 20x magnification. The most representative tissue block was selected for slide scanning.

#### Multiplexed immunofluorescence

##### Overview

We carried out multi-parameter quantification of epithelial and immune cell subsets and activation markers using the AkoyaBio Vectra Automated Imaging system at the MSKCC Parker Institute for Cancer Immunotherapy. We stained whole slides of formalin fixed paraffin embedded (FFPE) tissue for markers of ovarian cancer cells (panCK + CK8-CK18) and of specific leukocyte subsets, including macrophages (CD68), and cytotoxic T cells (CD8), known immune inhibitory proteins (PD-L1) and activation/exhaustion status of CD8 T cells (PD-1, TOX). Fields of view were chosen to include either entire tissue with minimal field overlap if the tissue was small, or a distribution of fields with 50% stroma/tumor at the edge plus some central areas of tumor dense fields. Marker intensities were QC’ed to fall in the range 5-30 a.u., and helped guide spectral unmixing. Lower values may be close to background and higher values prompted to check for channel spillage.

##### Tissue staining

Primary antibody staining conditions were optimized using standard immunohistochemical staining on the Leica Bond RX automated research stainer with DAB detection (Leica Bond Polymer Refine Detection DS9800). Using 4 µm formalin-fixed, paraffin-embedded tissue sections and serial antibody titrations, the optimal antibody concentration was determined followed by transition to a seven-color multiplex assay with equivalency. Optimal primary antibody stripping conditions between rounds in the seven-color assay were performed following 1 cycle of tyramide deposition followed by heat-induced stripping (see below) and subsequent chromogenic development (Leica Bond Polymer Regine Detection DS9800) with visual inspection for chromogenic product with a light microscope by a senior pathologist (T.J.H.). Multiplex assay antibodies and conditions are described in **Tab. S5**.

Tissue sections were baked for 3 hours at 62°C in vertical slide orientation with subsequent deparaffinization performed on the Leica Bond RX followed by 30 minutes of antigen retrieval with Leica Bond ER2 followed by 6 sequential cycles of staining with each round including a 30-minute combined block and primary antibody incubation (Akoya antibody diluent/block ARD1001).

For panCK and CK8-CK18, detection was performed using a secondary horseradish peroxidase (HRP)-conjugated polymer (Akoya Opal polymer HRP Ms + Rb ARH1001; 10-minute incubation). Detection of all other primary antibodies was performed using a goat anti-mouse Poly HRP secondary antibody or goat anti-rabbit Poly HRP secondary antibody (Invitrogen B40961/2; 10-minute incubation). The HRP-conjugated secondary antibody polymer was detected using fluorescent tyramide signal amplification using Opal dyes 520, 540, 570, 620, 650 and 690 (Akoya FP1487001KT, FP1494001KT, FP1488001KT, FP1495001KT, FP1496001KT, FP1497001KT).

The covalent tyramide reaction was followed by heat induced stripping of the primary/secondary antibody complex using Perkin Elmer AR9 buffer (AR900250ML) and Leica Bond ER2 (90% ER2 and 10% AR9) at 100°C for 20 minutes preceding the next cycle (1 cycle of stripping for CD68, PD1, PDL1, CD8, panCK/CK8/18 and two cycles for TOX). After 6 sequential rounds of staining, sections were stained with Hoechst (Invitrogen 33342) to visualize nuclei and mounted with ProLong Gold antifade reagent mounting medium (Invitrogen P36930).

##### Imaging and spectral unmixing

Seven color multiplex stained slides were imaged using the Vectra Multispectral Imaging System version 3 (Perkin Elmer). Scanning was performed at 20x (200x final magnification). Filter cubes used for multispectral imaging were DAPI, FITC, Cy3, Texas Red and Cy5. A spectral library containing the emitted spectral peaks of the fluorophores in this study was created using the Vectra image analysis software (Perkin Elmer). Using multispectral images from single-stained slides for each marker, the spectral library was used to separate each multispectral cube into individual components (spectral unmixing) allowing for identification of the seven marker channels of interest using Inform 2.4 image analysis software.

### Computational methods

#### Single-cell RNA sequencing

##### Overview

The pipeline is built using 10x Genomics *Martian* language and computational pipeline framework. *CellRanger* software (version 3.1.0) was used to perform read alignment, barcode filtering, and UMI quantification using the 10x GRCh38 transcriptome (version 3.0.0) for FASTQ inputs.

##### Quality control

CellRanger filtered matrices are loaded into individual *Seurat* objects using the *Seurat* R package (version 3.0.1) ^44,45^. The resulting gene by cell matrix is normalized and scaled for each sample. Cells retained for analysis had a minimum of 500 expressed genes and 1,000 UMI counts and less than 25% mitochondrial gene expression. Cell cycle phase was assigned using the *Seurat* CellCycleScoring function. *Scrublet* (version 0.2.1) was used to calculate and filter cells with a doublet score greater than 0.25. Sample matrices are merged by patient and subsequently renormalized and scaled using the default *Seurat* functions.

##### Major cell type identification

Major cell type assignments were computed on each patient with *CellAssign* (version 0.99.2) ^46^ using a set of curated marker genes. Marker genes were compiled for nine major cell types related to high-grade serous ovarian cancer (**Tab. S3**). These major cell types are defined as T cells, B cells, plasma cells, myeloid cells, dendritic cells, mast cells, endothelial cells, fibroblasts and ovarian cancer cells. Prior to running *CellAssign*, cells with zero expression for all marker genes were removed from the count matrix. Cell specific size factors are computed using *scran* (version 3.11). Default *CellAssign* parameters are used with a design matrix of patient batch labels. *CellAssign* returns a probability distribution over the major cell types and individual cells are labeled by the resulting most probable cell type.

##### Dimensionality reduction

Principal component analysis (PCA) was performed on the filtered feature by barcode matrix. UMAP embeddings including cohort-level and patient-level embeddings of all major cell types are based on the first 50 principal components. UMAP embeddings of major cell type supersets (see below) were based on the 50 batch-corrected harmony components. Diffusion map embeddings and pseudotime estimates were computed using R package *destiny* (v3.0.1) for the subset of CD8^+^ T cells (Angerer et al. 2015).

##### Batch correction and integration

Major cell types identified across samples were split into six supersets: (1) T cells; (2) B cells and plasma cells; (3) myeloid cells, dendritic cells and mast cells; (4) fibroblasts; (5) endothelial cells; (6) ovarian cancer cells. For each superset, R package *harmony* (version 0.1) was used for batch correction to account for patient-specific effects ^47^.

##### Clustering and cell subtype identification

Graph-based clustering is performed for each superset using the Louvain algorithm implemented in *Seurat* (version 3.0.1) at three different resolutions (0.1, 0.2, 0.3). Differential expression between identified clusters was computed using a Wilcoxon rank sum test as implemented in *Seurat* FindMarkers. Final results are filtered on log fold change > 0.25 and Benjamini-Hochberg adjusted *P*-value < 0.05. Clusters were annotated based on marker genes identified in differential gene expression analysis.

##### Gene signature scores

Cell state scores were calculated for the exhausted phenotype within the set of T cells using a manually curated list of genes as input to the *Seurat* AddModuleScore method ^48^. The curated list of genes was derived from a review of single-cell analyses of CD8^+^ T cell states in human cancers ^49^ (**Tab. S3**).

##### Patient specificity and TCGA subtyping

Patient specificity scores were computed with employing a shared nearest neighbour graph. For a given cell, patient specificity was defined as the observed fraction of nearest neighbors divided by the expected fraction of nearest neighbors in the patient subgraph. Here, the expected fraction of neighbors from the same patient was defined as the global fraction of cells for each patient. Scores were log_2_-transformed. Hence, a positive patient specificity score indicates an overrepresentation of cells derived from the same patient among its nearest neighbours, a negative score indicates an underrepresentation of same-patient cells and a score of 0 would reflect a perfectly mixed neighbourhood of patient labels. Consensus TCGA transcriptional subtypes were called using R package *consensusOV* (version 1.8.1) ^50^.

##### Generalized linear models of cluster composition

To estimate the effect of mutational signatures and tumor site specificity on the composition of cell clusters, we considered a generalized linear model (GLM) where we included interactions between signature, site and cluster identity for each major cell type defined in the scRNA, H&E and mpIF data. The data matrix includes the counts of every cluster *c* representing a cell type or cell state, sampled from site *s* in a patient with mutational signature subtype *m*. Using a binomial linear model one can analyze counts of repeated observations of cell types or cell states as binary choices,

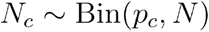

where *N_c_* is the count for cluster *c* in a sample, *N* is the total number of cells in the sample, and the probability to detect the cluster can be described by the logit function, 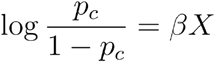.

To account for the effect of mutational signature and anatomic tumor site on cluster abundance observed in scRNA data, we formulate a GLM of the observed cell counts *N_c_* for a cluster described by the logit function, which is distributed as

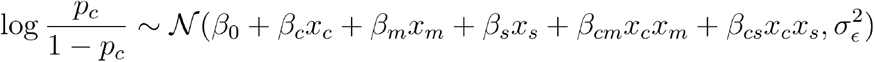

where *β*_0_ is a shared constant baseline per cluster that must be inferred, *β_c_*, *β_m_*, *β_s_* are individual fixed-effect terms to be inferred, *β_cm_* and *β_cs_* are cluster-signature and cluster-site interaction effects to be inferred, *x_c_*, *x_m,_* and *x_s_* are elements of the model design matrix *X*, and 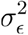 represents measurement noise. We note that for each cluster we have multiple measurement replicates of *N_c_* across signatures and sites. This formulation is used to fit a GLM of major cell types (**Fig. 2**), as well as to separately fit GLMs of cluster composition of cell states within each major cell type superset (cancer cells, **Fig. 4**; T cells, **Figs. 3,5**; myeloid cells, **Figs. 3,5**).

To model the abundance of major cell types in scRNA data from CD45^+^ and CD45^-^ samples, the GLM includes a covariate for CD45^+/-^ flow sorting with additional fixed-effect sorting coefficients *β_f_* and additional cluster-sorting interactions *β_cf_* to be inferred, plus an additional element *x_f_* in the model design matrix (**Fig. 2F**). Similarly, GLMs for H&E and mpIF data account for differences in cell type abundance observed in the tumor and stroma regions, incorporating a covariate for the tumor or stroma region counts with additional fixed-effect region coefficients *β_r_* and additional cluster-region coefficients *β_cr_* to be inferred, plus an additional element *x_r_* in the model design matrix (**Fig. 2F**).

#### Bulk whole genome sequencing

##### Alignment

Sequencing reads were aligned to the human genome reference GRCh37 (hg19) using the Burrows– Wheeler Aligner (BWA-MEM) v0.7.17-r1188 (https://sourceforge.net/projects/bio-bwa/).

##### SNVs and indels

Single nucleotide variant (SNV) and indels were called using *mutationSeq* (version 4.3.8; model v4.1.2.npz) available at https://github.com/shahcompbio/mutationseq. We also used *Strelka* (version 2.8.2) with default parameter settings to identify somatic SNVs and indels ^51^. Both SNVs and indels were then annotated for variant effects and gene-coding status using *SnpEff4* (version 5.0e). We identified a set of high confidence SNVs by taking the intersection of the high probability calls predicted from *mutationSeq* (with probability ≥ 0.9) and the somatic SNVs predicted from *Strelka*.

The high confidence set of SNVs were further filtered by removing the positions that fell within either of the following regions: (i) the UCSC Genome Browser blacklists (Duke and DAC), and (ii) defined in the ‘CRG Alignability 36mer track’ with more than two nucleotide mismatches, requiring a 36-nucleotide fragment to be unique in the genome even after allowing for two differing nucleotides. Post processing on this set of high confidence SNVs and somatic indels from *Strelka* involved removing the known variants (both SNVs and indels) that were obtained from the 1000 Genomes Project (release 20130502) and dbSNP (version dbsnp 142.human 9606). The set of high confidence somatic SNVs and indels passing the above filters were then used in feature computation for mutational signature analysis, and high confidence somatic SNVs were also used for neoantigen prediction.

##### Rearrangements

Rearrangement breakpoints were predicted using *lumpy* (version 0.2.12) ^52^ executed by *SpeedSeq* version 0.1.08 ^53^, and *destruct* (version 0.4.18) derived from *nFuse* ^54^, available at https://github.com/amcpherson/destruct. In brief, *destruct* extracted discordant and non-mapping reads from BAM files and realigned the reads using a seed-and-extend strategy. Split alignment across a putative breakpoint was attempted for reads that did not fully align to a single locus. Discordant alignments were clustered according to the likelihood they were produced from the same breakpoint. Multiple mapped reads were assigned to a single mapping location using previously described methods ^55^. Finally, heuristic filters removed predicted breakpoints with poor discordant read coverage of sequence flanking predicted breakpoints.

We applied a stringent 3-step filtering criteria to identify high confidence breakpoint calls for downstream analysis, as follows:

Step 1: Breakpoints that were predicted by both algorithms, *lumpy* and *destruct*, were taken.
Step 2: We removed (i) the breakpoints from the poor mappability regions, (ii) events with break distance ≤30 bp, (iii) breakpoints annotated as deletion with breakpoints size <1,000 bp. Furthermore, only high confidence breakpoints that had at least five supporting reads in tumor and no read support in the matched normal sample were used in the analysis. The breakpoints were further filtered by removing the positions in either of the following regions: (i) UCSC Genome Browser blacklists (Duke and DAC), and (ii) defined in the ‘CRG Alignability 36mer track’ with more than two nucleotide mismatches, requiring a 36-nucleotide fragment to be unique in the genome even after allowing for two differing nucleotides.
Step 3: Predictions with small break distance and low number of supporting reads in tumor samples were excluded.

#### Myriad HRD test

We used a commercial assay (Myriad Genetics ‘myChoice CDx’) to test for genome-wide LOH, the number of chromosomal breakpoints in large scale state transitions and telomeric allelic imbalance. If the resulting HRD score is greater than 42 the sample was deemed HR-deficient.

#### Targeted sequencing (MSK-IMPACT)

Genomic DNA isolated from FFPE tumor tissue and matched normal blood was subjected to hybridization capture and sequenced with deep coverage (700x) ^56^. Variant calling for the MSK-IMPACT gene panel and copy number analysis was performed using the MSK-IMPACT clinical pipeline (https://github.com/mskcc/Innovation-IMPACT-Pipeline).

#### Mutational signatures

We analyzed mutational signatures by integrating point mutations and structural variations detected by bulk whole genome sequencing in a unified probabilistic approach called multi-modal correlated topic models (MMCTM) ^11^. MMCTM analysis enables robust determination of mutational signatures, their correlation structure and the delineation of sub-groupings on the basis of point mutation signatures ^57^ and structural variations.

We estimated signature probabilities for bulk WGS samples in the SPECTRUM cohort (n=21) using the MMCTM, based on SNV and SV signatures inferred from HGSOC (n=170) and triple-negative breast cancer (n=139) bulk whole genomes (total n=309) (**Fig. S4A**). By clustering the SPECTRUM cohort samples together with the 309 HGSOC and TNBC samples using UMAP and HDBSCAN ^58^, we assigned the 21 SPECTRUM samples into one of 9 strata defined solely by SNV and SV signature probabilities. These strata include those with samples enriched for: i) *BRCA1*-associated homologous recombination deficient (HRD) point mutation signatures accompanied by tandem duplications (HRD-Dup), ii) *BRCA2*-associated HRD point mutation signatures accompanied by interstitial deletions (HRD-Del), iii) *CDK12*-associated tandem duplications (TD) and iv) foldback-inversions mediated by the breakage-fusion bridge process (FBI) (**Fig. S4A**). These strata are associated with distinct prognostic profiles under standard of care treatment ^12^.

Mutational signatures for cases without bulk WGS data were assigned based on gene-level mutations in MSK-IMPACT, based on the presence of *BRCA1* (HRD-Dup), *BRCA2* (HRD-Del) and *CDK12* (TD) loss-of-function mutations or homozygous deletions, and *CCNE1* amplifications (FBI). Additionally, cases with Myriad Genetics ‘myChoice CDx’ data were labelled as HRD-Other based on a positive test score.

Consensus mutational signatures were preferentially derived based on: i) MMCTM signatures derived from bulk WGS, ii) gene-level annotations in MSK-IMPACT, and iii) Myriad ‘myChoice CDx’ test results. Mutational signatures for two cases without other informative data (patients 037 and 112) were resolved based on single-cell whole genome sequencing.

#### Neoantigen prediction

To predict the peptide binding affinity of neoantigens *in silico* using tumor and matched normal WGS samples (n=21), candidate nonsynonymous SNVs were used to generate a list of peptides of amino acid length 8, 9, 10 and 11. The binding affinity of each mutant peptide and its corresponding wildtype peptide to the patient’s germline HLA alleles were predicted using *NetMHCpan* v4.1 in *pVACtools*^64^. Peptides with inferred mutant binding affinities below 1,000 nM are defined as neoantigens.

#### HLA loss of heterozygosity

To detect allele-specific copy number loss of heterozygosity (LOH) of the HLA locus in single cells profiled by scRNA-seq, we inferred allele-specific alterations in chromosome arm 6p which harbors HLA class I and II genes using *schnapps* ^59^. We first called germline heterozygous SNPs in the scRNA-seq tumor data using *cellSNP* ^60^. As input, we used the set of heterozygous SNPs identified in the corresponding normal WGS dataset for each sample. The liftover script provided in *cellSNP* was used to lift over SNP coordinates from the GRCh37 (hg19) to the GRCh38 reference genome. Following genotyping, we aggregated SNP counts across all cells and defined the B allele as the allele with lowest allele frequency for each SNP. As SNP counts are very sparse in scRNA-seq, we then aggregated cell-level counts of the B-allele across chromosome arms in order to compute the BAF for each arm in each cell. We then generated a cell by chromosome arm BAF matrix and incorporated this into the Seurat gene expression objects. To assign allelic imbalance states (balanced, imbalanced, LOH) to chromosome arms in each cell we used the mean BAF of each arm per cell as follows: balanced (BAF ≥ 0.35); imbalanced (0.15 ≤ BAF < 0.35); LOH (BAF < 0.15). Documentation and code is available at https://shahcompbio.github.io/schnapps/.

To validate our observations of allele-specific alterations in chromosome arm 6p in relation to the HLA locus, we detected gene-level HLA class I LOH from tumor and matched normal MSK-IMPACT data using *LOHHLA* (McGranahan et al., 2017). Tumor samples from 1,111 cases in the MSK-IMPACT cohort with HGSOC histology were selected, based on a HGSOC or HGSFT OncoTree classification ^61^. This cohort is a superset that includes samples from MSK SPECTRUM patients. Patient HLA references were built from tumor and normal MSK-IMPACT reads using *Polysolver* v4^62^. Tumor purity and ploidy were estimated using *FACETS* ^63^ and used for subsequent HLA LOH analysis. HLA LOH was called for an allele in the tumor sample using *LOHHLA*. LOH was observed for each HLA gene if the estimated copy number was < 0.2 and the significance of allelic imbalance was *P* < 0.01, which tests for pairwise differences in logR values between the two HLA homologs (paired *t*-test).

#### Digital histopathology

We built a training dataset of cellular annotations of scanned H&E images. Expert delineation and quantification of cell and tissue types present in the H&E slides was carried out on MSK Slide Viewer, a computational pathology interface for review and annotation of histopathology images. Nuclear segmentation was carried out using StarDist, a method for nuclear detection based on the U-Net neural network architecture ^65,66^. Membrane segmentation was approximated using a cell expansion of 3 μm of the nuclear boundary. The training dataset encompasses a set of 61 slides from a representative set of patients and sites. To classify regions of tumor, stroma, vasculature and necrosis, we trained an ANN-based pixel classifier using QuPath v0.2.3 ^65^, which operates on higher-order pixel features over multiple channels and scales within the image. In addition, lymphocytes and “other” cells were annotated in 19 of these slides by a researcher using MSK Slide Viewer. After importing these annotations into QuPath, along with cellular segmentations and feature vectors generated from StarDist, we then trained an ANN-based cellular classifier which operates over cellular measurements to identify lymphocytes. We then applied these models for inference across whole-slide H&E images over the larger cohort, and used these model outputs to compute statistics on lymphocytic densities and other spatially-derived measurements.

#### Multiplexed immunofluorescence

We carried out nuclear segmentation based on DAPI intensity using the watershed algorithm in QuPath v0.2.3 ^65^, setting a minimum DAPI threshold of 1 a.u. with an expected nucleus area ranging between 5 μm^2^ and 100 μm^2^. Membrane segmentation was approximated using a cell expansion of 3 μm of the nuclear boundary. Starting from 1,194 quality-filtered FOVs across 90 tissue samples from 35 patients, segmentation yielded a total of 9,257,609 cells. To annotate regions of tumor and stroma, we trained a pixel classifier with examples of panCK^+^ (tumor) and panCK^-^ regions (stroma). Following nuclear segmentation, we extracted the pixel intensities per cell for functional markers expressed in the cytoplasm (panCK, CD68, CD8, PD1, PDL1) and in the nucleus (TOX) in order to define cell types and cell states. All channels were manually thresholded in at least one field of view (FOV) per slide, and marker positivity was determined by setting these thresholds on the mean pixel intensity. Segmented objects which were double- or triple-positive for multiple cell type markers (panCK, CD68, CD8) were counted as separate cells, yielding a total of 10,663,919 single cells. Marker assignments were used to define cell states of epithelial cells (panCK^+^PDL1^-^, panCK^+^PDL1^+^), macrophages (CD68^+^PDL1^-^, CD68^+^PDL1^+^) and CD8^+^ T cells (CD8^+^PD1^-^TOX^-^, CD8^+^PD1^+^TOX^-^, CD8^+^PD1^+^TOX^+^).

Analysis of spatial topology comprised estimation of spatial densities and inter-cellular nearest-neighbor distances. Spatial density estimates as a function of distance to the tumor-stroma boundary were obtained by aggregating cell counts within 10 μm distance bands from the boundary in each FOV, grouped across FOVs, and normalized by the total number of cells for a given phenotype of interest. Error bars were calculated as the standard error of the probability *p* to observe a given phenotype as 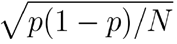, where *N* is the total number of cells in that distance band. Inter-cellular distances between nearest neighbors are calculated using the distance matrix *r_ij_* between cells *i* and *j*, where the value of the (*i*, *j*) element in the matrix is the radial distance from cell *i* to cell *j*. Once the per-cell nearest neighbors have been computed, the summary statistics over nearest neighbor distances per phenotype can be estimated. Proximity counts between phenotypes within a fixed radius *R* can also be determined based on the per-cell nearest neighbors.

---

**Figure S1.**
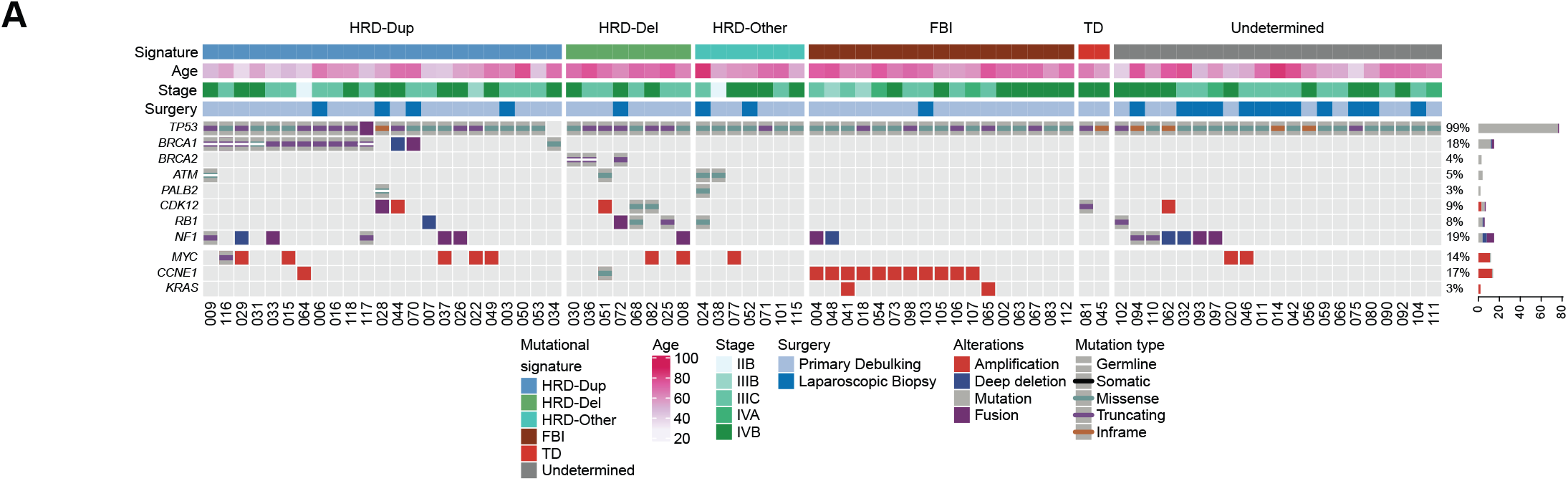
Prevalence of germline and somatic mutations, related to Fig. 1. **A)** Recurrent alterations in oncogenes and tumor-suppressor genes in the MSK SPECTRUM cohort, detected by MSK-IMPACT. Bars indicate the percentage of cases harboring different classes of genomic alterations. Mutation types are broken down into missense variants, truncating variants (nonsense mutations, frameshift indels, splice site variants) and inframe variants. Mutations shown are somatic except those highlighted as germline variants. Labels on the top indicate the mutational signature subtype, patient age, staging, and type of surgical procedure. Patients were staged following FIGO Ovarian Cancer Staging guidelines. Patients in the neoadjuvant setting were given a clinical stage based on pre-treatment imaging, and patients that received up-front surgery (staging or primary debulking) were given a pathological stage based on the specimens collected.

**Figure S2.**
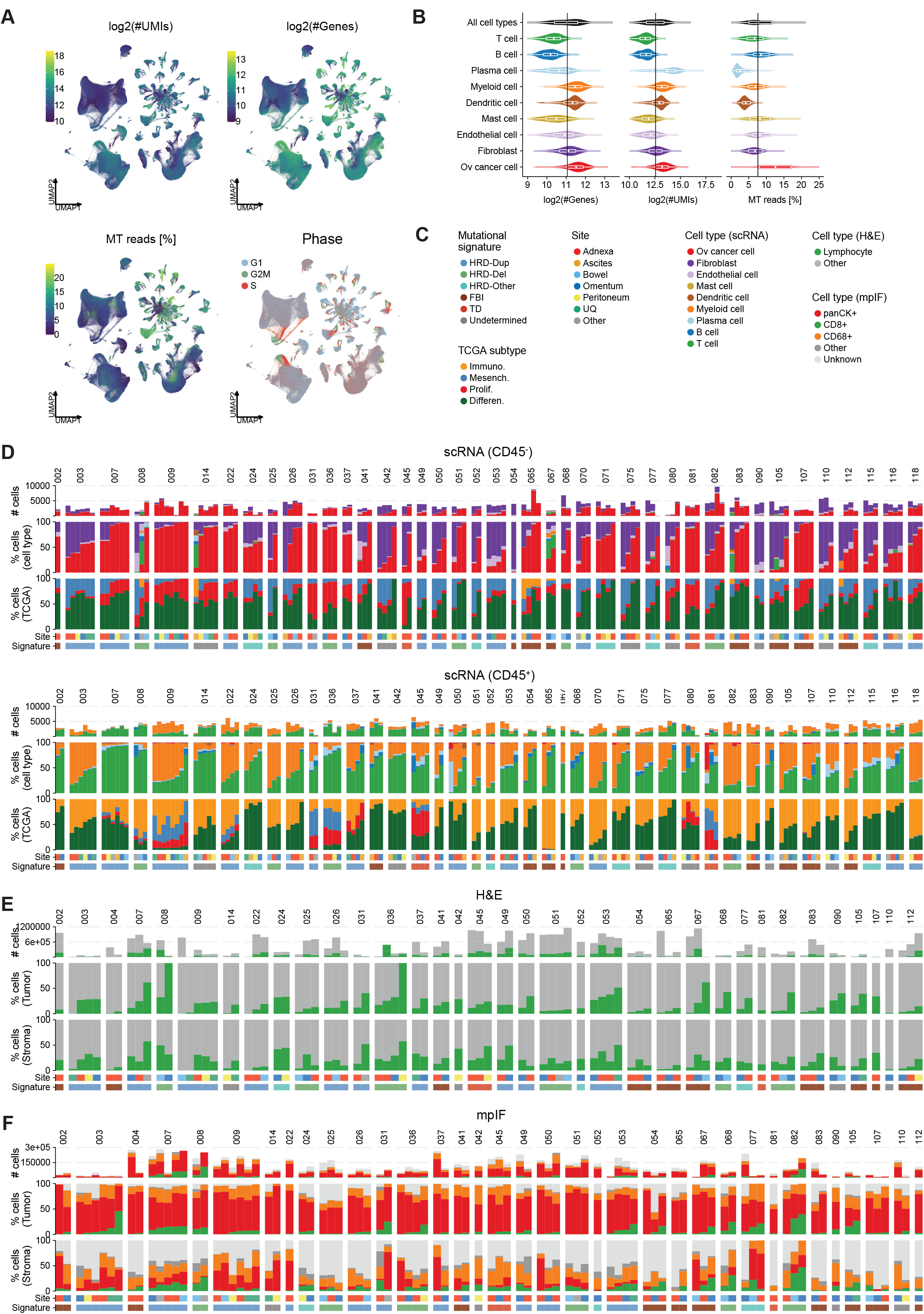
Quality control of scRNA-seq data and cell type abundance profiled by scRNA-seq, H&E and mpIF, related to Fig. 2. **A)** UMAPs of cells profiled by scRNA-seq colored by different QC metrics: log_2_ transformed number of UMIs and genes, fraction of mitochondrial reads, cell cycle phase. **B)** Distributions of QC metrics per cell type. **C)** Color legend for D)-F). **D)** Absolute and relative cell type compositions and TCGA subtype compositions of CD45^-^ (top) and CD45^+^ (bottom) sorted samples based on scRNA, separated by patient, ranked by fraction of ovarian cancer cells and T cells respectively. **E)** Absolute and relative cell type compositions based on H&E, ranked by lymphocyte fractions for tumor-rich (top) and stroma-rich (bottom) compartments. Panels analogous to D). **F)** Absolute and relative cell type compositions based on mpIF, ranked by CD8^+^ T cell fractions in tumor-rich (top) and stroma-rich (bottom) compartments. Panels analogous to D).

**Figure S3.**
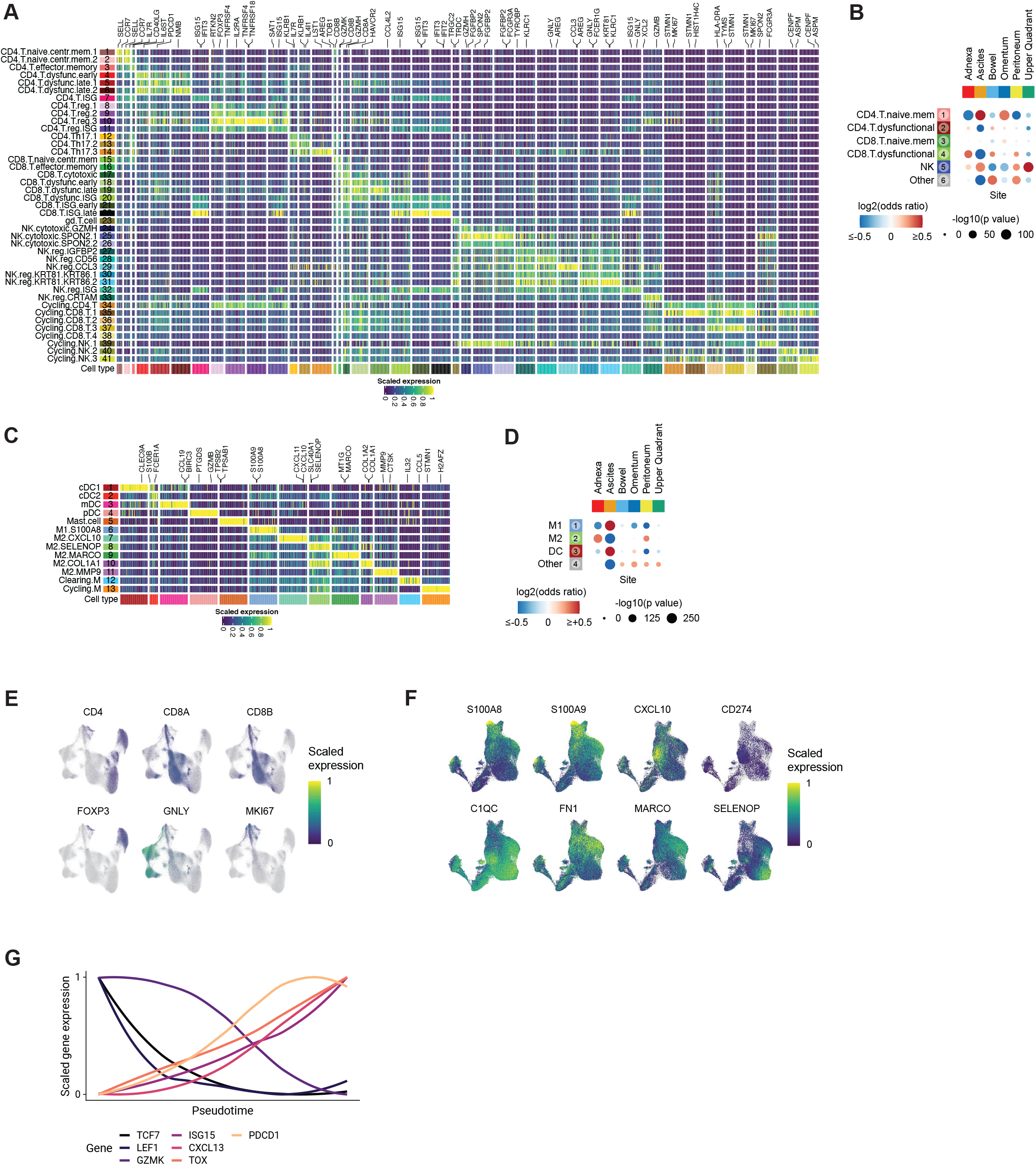
Anatomic site specificity and marker gene expression of T, NK and cell myeloid phenotypes, related to Fig. 3. **A)** Heatmap of scaled marker gene expression (averaged per cluster) for T and NK cell clusters, showing differentially expressed genes in columns and clusters in rows. Genes are grouped by cluster. Top 2 genes per cluster are highlighted. **B)** Site-specific enrichment of coarse-grained T/NK cell clusters using GLM. Color gradient indicates log_2_ odds ratios and sizes indicate the BH-corrected -log_10_(p value). **C)** Marker gene expression heatmap for myeloid cells (dendritic cells, mast cells and macrophage clusters). **D)** Site-specific enrichment of coarse-grained myeloid cell clusters using GLM analogous to B). **E)** UMAPs of T and NK cells showing scaled expression of marker genes of interest. **F)** UMAPs of macrophage cells showing scaled expression of marker genes of interest. **G)** Genes of interest in subsets of CD8^+^ T cells along pseudotime axis.

**Figure S4.**
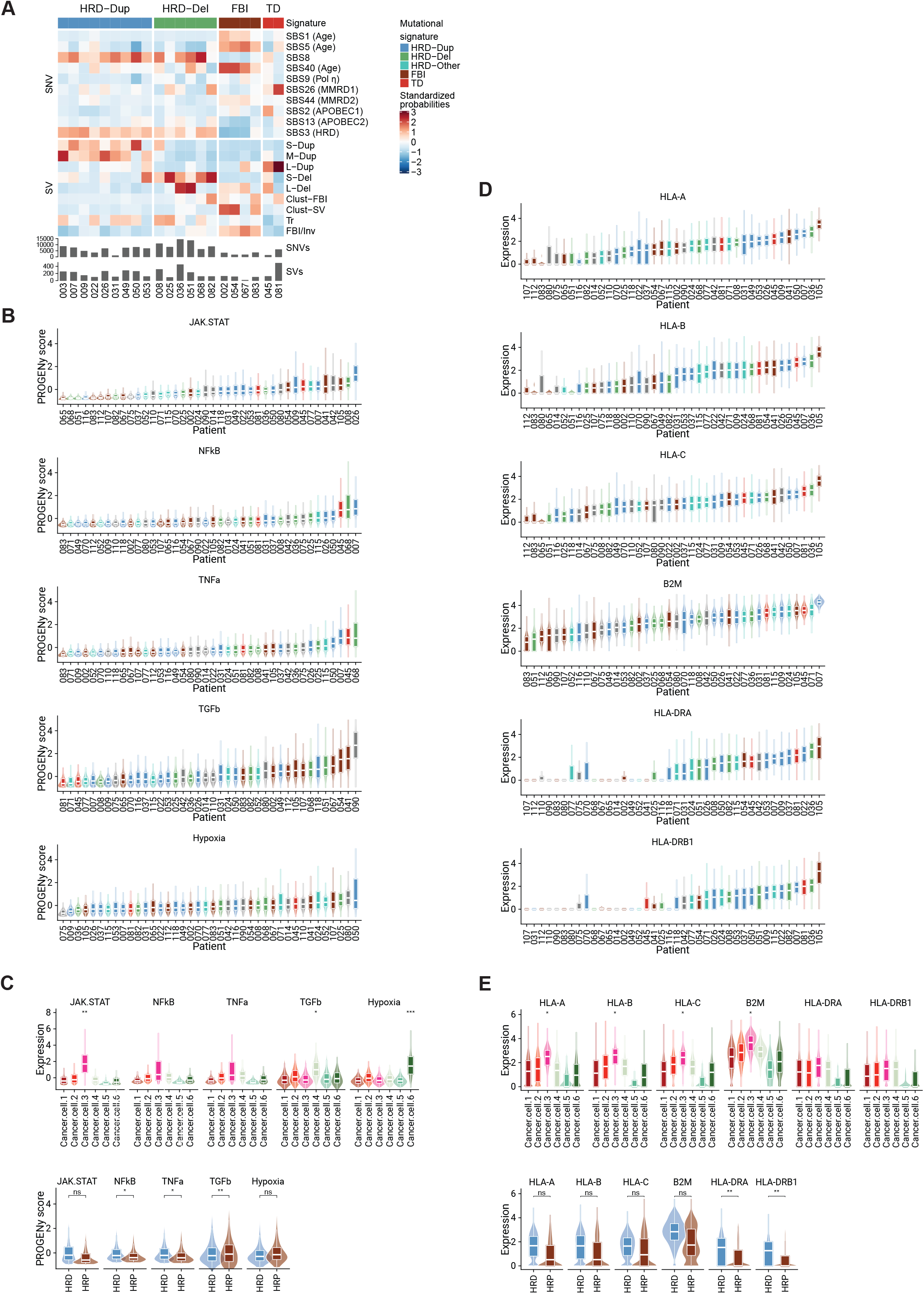
Mutational signatures and their impact on cancer cell-intrinsic signaling, related to Fig. 4. **A)** Heatmap of standardized probabilities for genomic features used to infer mutational signature subtypes from whole genome sequencing. Patients (in columns) are grouped by mutational signature. Features used for inference (in rows) are grouped into single nucleotide variant (SNV) and structural variation (SV) features. SV features include duplications (S-Dup, M-Dup, L-Dup), deletions (S-Del, L-Del), unclustered and clustered foldback inversions (FBI/Inv, Clust-FBI), clustered rearrangements (Clust-SV) and translocations (Tr). Bar graphs indicate total number of SNVs and SVs per tumor sample. **B)** Single cell distributions of PROGENy pathway activity per patient. **C)** Single cell distributions of PROGENy pathway activity per cluster (top subpanel) and HR status (bottom subpanel). **D)** Single cell distributions of HLA class I and class II gene expression per patient. **E)** Single cell distributions of HLA gene expression per cluster (top subpanel) and HR status (bottom subpanel). \**P*<0.05, \*\**P*<0.01, \*\*\**P*<0.001, \*\*\*\**P*<0.0001. Brackets: Wilcoxon pairwise comparisons.

**Figure S5.**
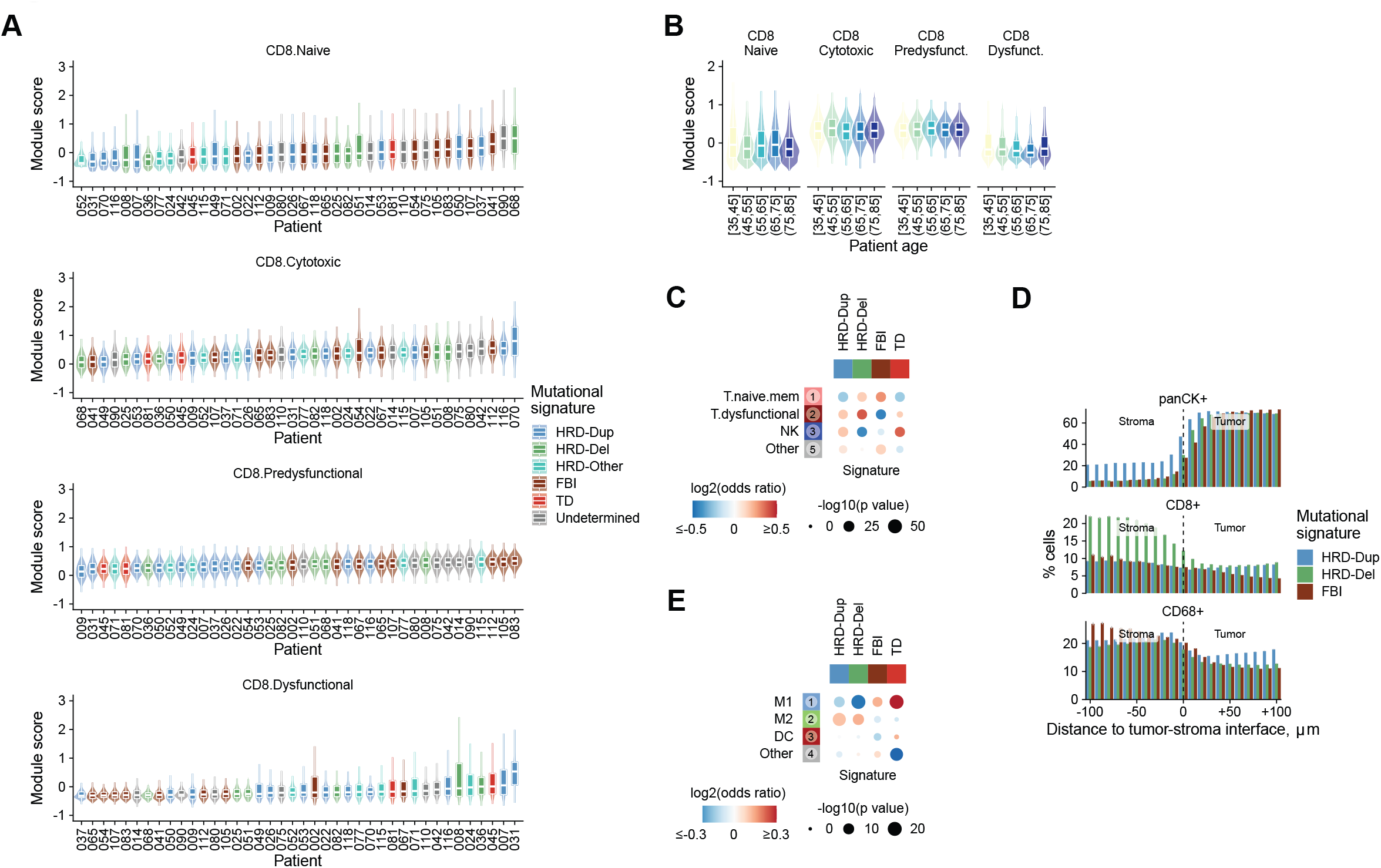
HR deficiency and tumor immunogenicity impact T cell phenotypes, related to Fig. 5. **A)** Single cell distributions of T cell module scores per patient. **B)** Single cell distributions of T cell module scores as a function of patient age groups. **C)** Signature-specific enrichment of coarse-grained T/NK cell clusters using GLM. Color gradient indicates log_2_ odds ratios and sizes indicate the BH-corrected -log_10_(p value). **D)** Spatial density of cancer cells, CD8^+^ T cells and macrophages as a function of distance to the tumor-stroma interface, grouped by mutational signature. Counts within 10 μm distance bands are grouped across FOVs from each mutational signature subtype, and are normalized by the total number of cells. **E)** Signature-specific enrichment of coarse-grained myeloid cell clusters using GLM analogous to C). \**P*<0.05. Brackets: Wilcoxon pairwise comparisons.

**Figure S6.**
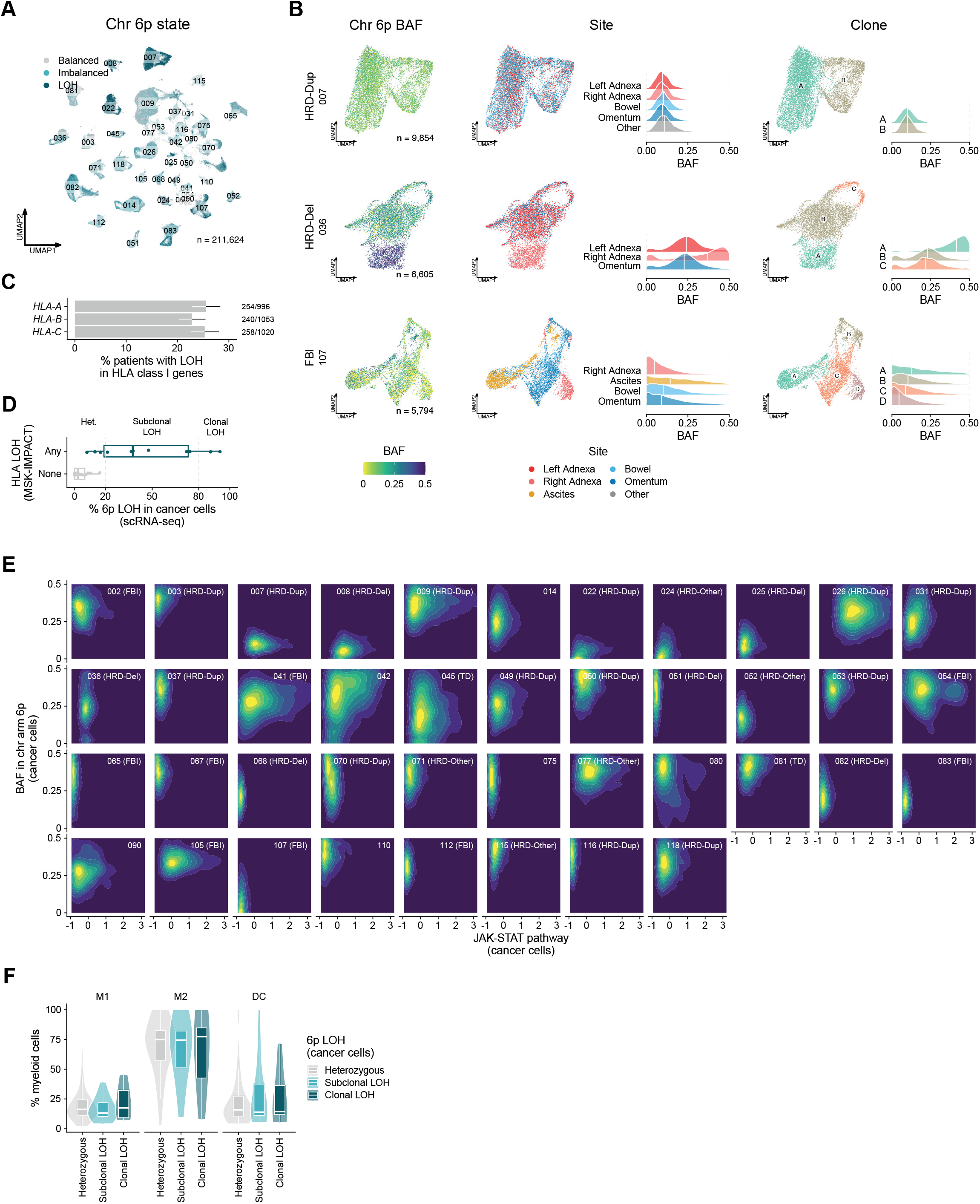
Intratumor heterogeneity of HLA loss of heterozygosity and its impact on immune phenotypes, related to Fig. 6. **A)** Allelic state of chromosome arm 6p. Allelic imbalance states per cell are assigned based on the mean 6p BAF per cell as balanced (BAF ≥ 0.35), imbalanced (0.15 ≤ BAF < 0.35) or LOH (BAF < 0.15) (**Methods**). **B)** UMAPs of cancer cells profiled by scRNA-seq from representative patients of each mutational subtype, highlighting intratumor heterogeneity in 6p B-allele frequency (BAF). From left to right, UMAPs are colored by 6p BAF, tumor site and tumor clone. Clones are defined using patient-level Louvain clustering of cancer cells. Density plots show site-specific and clone-specific 6p BAF distributions. Only sites with ≥10 cancer cells are shown. **C)** Percentage of patients with LOH of HLA class I genes in the MSK-IMPACT HGSOC cohort (n=1,111 patients). **D)** Validation of median 6p BAF estimates in cancer cells profiled by scRNA-seq using HLA LOH status in site-matched MSK-IMPACT samples. 27 out of 41 patients profiled by scRNA-seq have site-matched MSK-IMPACT data. **E)** Normalized density contours of 6p BAF and JAK-STAT pathway activity in cancer cells for each patient. **F)** Fraction of M1 macrophages, M2 macrophages and dendritic cells as a function of clonality of 6p LOH in cancer cells. Percentage allelic loss of chromosome 6p arm in cancer cells per site is used to bin samples according to their 6p LOH status (heterozygous: % 6p LOH < 20%, clonal LOH: % 6p LOH > 80%). In panels D)-F), only BAF estimates from cells with ≥10 reads aligning to chromosome arm 6p are considered, and allelic imbalance states are assigned per cell based on the mean 6p BAF per cell as balanced (BAF ≥ 0.35), imbalanced (0.15 ≤ BAF < 0.35) or LOH (BAF < 0.15) (**Methods**).

**Figure S7.**
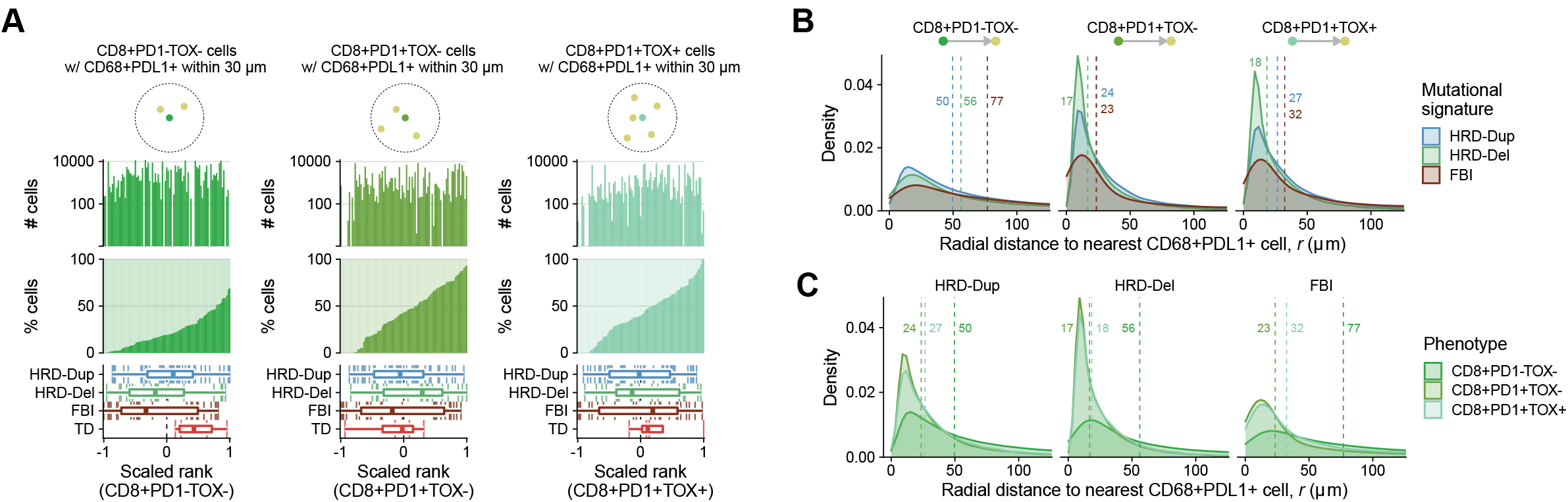
Spatial topology of interactions between T cells and macrophages, related to Fig. 7. **A)** Proximity analysis between CD8^+^ T cell phenotypes and CD68^+^PDL1^+^ macrophages based on mpIF data, ranking samples by the fraction of CD8^+^PD1^-^TOX^-^ T cells (left), CD8^+^PD1^+^TOX^-^ T cells (middle) or fraction of CD8^+^PD1^+^TOX^+^ T cells (right) with ≥1 CD68^+^PDL1^+^ cell within 30 μm. Vertically aligned subpanels share the same x-axis. Upper panels: Bar graphs show absolute abundance of CD8^+^ T cell states. Middle panels: Bar graphs show the fraction of CD8^+^ T cell phenotypes with ≥1 CD68^+^PDL1^+^ cell within 30 μm. Bottom panels: Box plot distributions of sample ranks with respect to mutational signature. **B and C)** Nearest-neighbor distance from CD8^+^ T cell phenotypes to CD68^+^PDL1^+^ macrophages aggregated across fields of view, grouped by mutational signature subtype. Vertical lines indicate the median nearest-neighbor distance.

## TABLES

**Table S1** Clinical overview of the MSK SPECTRUM patient cohort. Related to Figure 1.

**Table S2** Sample inventory. Metadata associated with scRNA-seq, H&E, mpIF, bulk tumor and normal WGS, Myriad HRD tests, and tumor and normal MSK-IMPACT datasets. Related to Figure 1.

**Table S3** Cell type and cell subtype markers. Clusters are annotated based on marker genes identified in differential gene expression analysis. Related to Figures 2-7.

**Table S4** Mutational signature proportions and mutational subtype assignments from WGS datasets for SPECTRUM patients. Related to Figures 1-7.

**Table S5** Antibodies and staining conditions for multiplexed immunofluorescence. Related to Figures 2 and 7.

